# A commensal-derived sugar protects against obesity by regulating immunometabolism

**DOI:** 10.1101/2024.06.12.598703

**Authors:** Chin Yee Tan, Yinghui Li, Danting Jiang, Barbara S. Theriot, Meghana V. Rao, Neeraj K. Surana

## Abstract

Obesity is a worsening global epidemic that is regulated by the microbiota through unknown bacterial factors. We discovered a human commensal bacterium, *Clostridium immunis*, that treats obesity by secreting a phosphocholine-modified exopolysaccharide. Loss- and gain-of-function bacterial mutants involving the phosphocholine biosynthesis locus (*licABC*) revealed the phosphocholine moiety is critically required to protect against metabolic disease. This *C. immunis* exopolysaccharide decreases small-intestinal and visceral fat levels of IL-22, which increases metabolic activity specifically in visceral adipose tissue. Importantly, phosphocholine biosynthesis genes are less abundant in humans with obesity or hypertriglyceridemia, findings that suggest the role of bacterial phosphocholine is conserved across mice and humans. These results define a bacterial molecule—and its key structural motif—that provides immunometabolic control of obesity. More broadly, they highlight a clinically translatable strategy to reduce visceral fat.

## INTRODUCTION

The global obesity epidemic continues to worsen, with over half the world’s population estimated to be overweight or obese by 2035^1^. This increase in the prevalence of obesity is estimated to drive sharp increases in cardiometabolic diseases^2,3^, such as coronary heart disease, type 2 diabetes, and stroke. Although body mass index (BMI) is frequently used as a diagnostic tool for obesity, it is well documented to be an imperfect measure^4–7^. Instead, excess visceral adipose tissue (VAT)—the hormonally active component of body fat that surrounds the internal organs^8^—is a better predictor for development of metabolic disease^8–11^, with many suggesting it be used to diagnose obesity^12^.

While the development of visceral adiposity is complex, it is increasingly recognized that the microbiota plays a critical role in its regulation. Obese and lean humans have marked differences in the composition and functional potential of their gut microbiomes^13–15^, but it has been challenging to disentangle the cause–effect relationship between the microbiome and obesity in human studies^16^. Mendelian randomization studies and host genetics-based approaches have linked specific bacterial taxa (e.g., *Christensenella*, *Akkermansia*, and *Bifidobacterium* spp.) and metabolites (e.g., short-chain fatty acids, bile acids) to changes in metabolic health^17–20^, with the directionality of the effect dependent on the specific feature. Although microbiome-based intervention studies have thus far been largely unsuccessful in treating obesity in humans^21^, animal studies have convincingly demonstrated that the microbiome causally regulates adiposity and weight. These effects stem, in part, from the microbiota harvesting greater amounts of energy from the host diet, promoting storage of that energy as fat, and regulating the quality of the adipose tissue^22–24^. However, the specific bacterial products and host mechanisms underlying host–microbiome interactions that modulate VAT and metabolic diseases remain poorly understood.

Here, we demonstrate that *Clostridium immunis*, a human commensal bacterium we previously found protects against colitis^25^, also protects against obesity, with decreases in weight gain, serum triglyceride levels, and VAT. Importantly, we purified and characterized an exopolysaccharide (EPS) as the bioactive molecule required for these metabolic effects, defined the EPS structural motif critical for activity, elucidated the immunological mechanism of action, and identified its metabolic effects that lead to disease protection. Moreover, we used metagenomic analyses to confirm our key findings are generalizable to metabolic disease in humans. Considered together, our study provides new mechanistic insight into how the microbiome regulates VAT and metabolic disease.

## RESULTS

### *C. immunis* protects against obesity

In previously published work, we identified *Clostridium immunis*, a newly-discovered human commensal bacterium that protects mice against colitis^25^. To better understand how *C. immunis* protects against colitis, we compared the colonic transcriptional profile of gnotobiotic mice harboring a mouse microbiota (MMb) treated with or without *C. immunis*. Although we did not detect notable changes in immune-related genes, we observed that *C. immunis* treatment surprisingly led to suppression of numerous genes related to lipid uptake and metabolism (Figure S1A). We confirmed these findings extended to the small intestine, the primary site of lipid metabolism, by demonstrating *C. immunis* inhibits jejunal expression of *Cd36*, which encodes a transporter that imports fatty acids into cells^26^, and *Scd1*, which encodes a stearoyl-coenzyme A desaturase^27^ (Figures S1B, S1C). Moreover, we found that *C. immunis* treatment inhibits small-intestinal expression and protein levels of *Nfil3* (Figures 1A, S1D), a microbiota-dependent transcription factor that positively regulates the expression of many of the differentially expressed metabolism-related genes we identified (Figure S1E)^28^.

**Figure 1:**
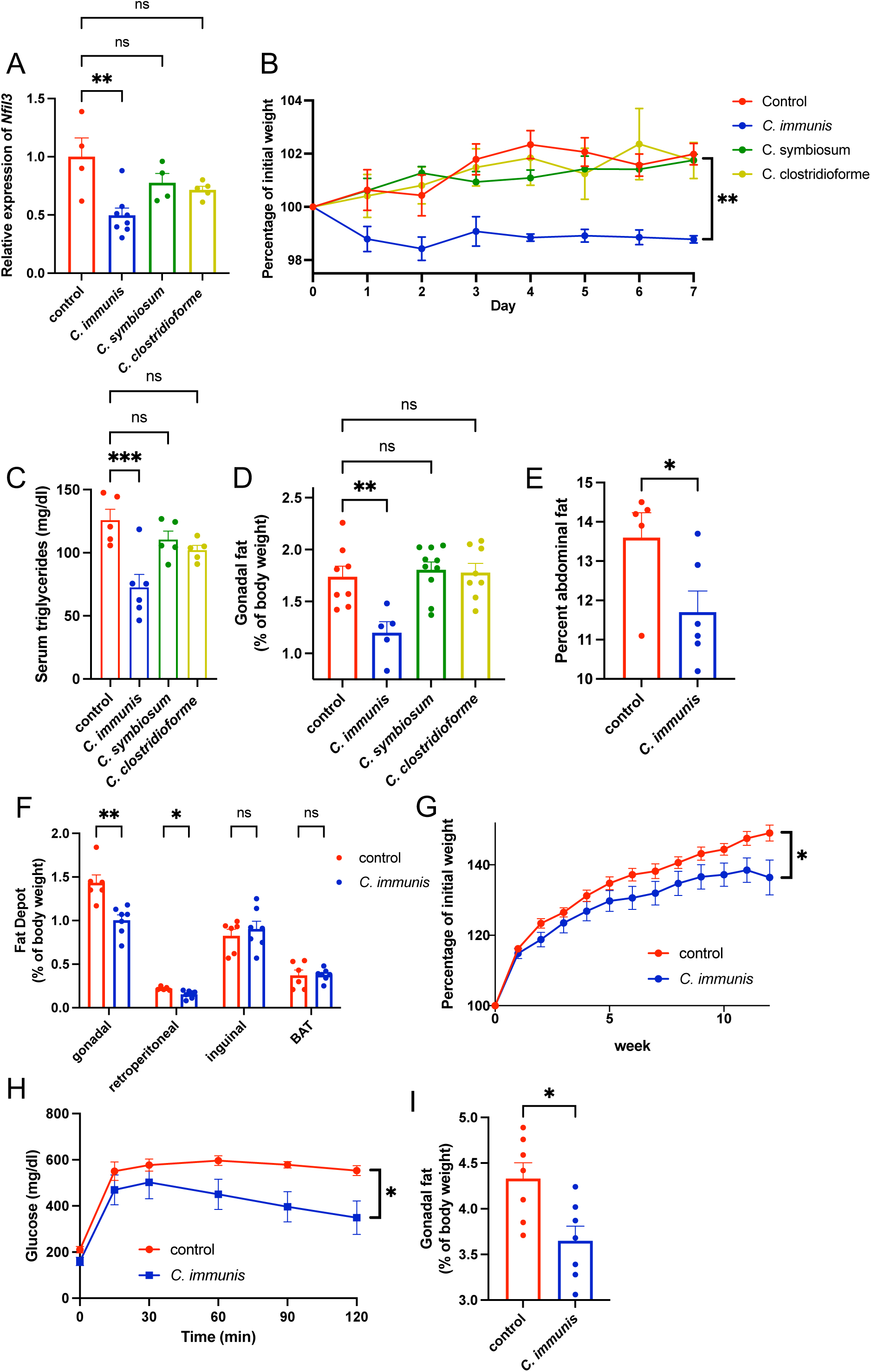
*C. immunis* protects against obesity. (**A**) Quantitative PCR (qPCR) analysis of small-intestinal *Nfil3* expression from gnotobiotic C57BL/6 mice harboring a mouse microbiota (MMb) treated orally with or without the indicated bacteria. (**B–F**) SPF C57BL/6 mice were orally treated with or without the indicated bacteria. Body weight was assessed daily for one week (**B**; *n* = 5 mice per group). One week after bacterial treatments, serum triglycerides (**C**), gonadal fat mass normalized to body weight (**D**), percent abdominal fat measured via a DEXA scan (**E**), and the mass of different fat depots normalized to body weight were assessed (**F**). (**G–I**) Longitudinal weights for SPF C57BL/6 mice fed a high-fat diet for 12 weeks and orally treated with or without *C. immunis* each week (**G**; *n* = 7 mice per group). Serum glucose values from an intraperitoneal glucose tolerance test conducted after the 12 weeks of high-fat diet are depicted (**H**), and gonadal fat mass normalized to body mass was assessed (**I**). Graphs represent means±s.e.m. \**P* < 0.05; \*\**P* < 0.01; \*\*\**P* < 0.001; ns, not significant by one-way ANOVA with Holm-Šidák correction (**A–D**) or Student’s t-test (**E–I**). For panels B, G, and H, statistical comparisons were performed on the area under the curve (AUC), which used background-corrected glucose values for panel H. See also Figure S1.

Given that deletion of any of these three genes results in decreased lipid metabolism and reduced VAT^27–29^, we reasoned that treatment with *C. immunis* may similarly improve metabolic features. Indeed, one week after a single oral treatment with *C. immunis*, specific pathogen-free (SPF) C57BL/6 mice had a decrease of ∼3% in body weight, ∼25% in serum triglyceride levels, and ∼33% in gonadal white adipose tissue (gWAT; a measure of VAT) compared to untreated mice (Figures 1B–D, S1F–H). We additionally confirmed *C. immunis* reduced abdominal adiposity via dual energy X-ray absorptiometry (DEXA; Figure 1E), with no difference in lean mass (Figure S1I). Surprisingly, this reduction in adiposity was specific to VAT as both gWAT and retroperitoneal fat depots were decreased by ∼33%, with no impact on subcutaneous inguinal white adipose tissue (iWAT) or interscapular brown adipose tissue (BAT; Figure 1F).

While these results demonstrate that *C. immunis* improves metabolic disease under short-term, healthy conditions, we investigated whether it also protects against these features in a model of diet-induced obesity (DIO). We orally treated mice with or without *C. immunis* every week for 12 weeks, during which time mice were fed a high-fat diet. While control mice gained ∼50% of their body weight during this time, *C. immunis*-treated mice gained ∼13% less weight (Figures 1G, S1J). In addition, *C. immunis*-treated mice had improved glucose tolerance and significantly less gWAT at the conclusion of the high-fat diet challenge (Figures 1H, 1I). These data demonstrate that *C. immunis* is also effective at reducing weight gain and adiposity in the context of long-term DIO. Although the microbiota generally promotes adiposity^28,30^, it is notable that treatment with *C. immunis* decreased VAT, even in the presence of a complex microbiota.

### A high-molecular weight exopolysaccharide purified from *C. immunis* recapitulates its activity

Intriguingly, neither *Clostridium symbiosum* nor *Clostridium clostridioforme*, two bacterial species that share significant genetic and proteomic similarity with *C. immunis*, respectively^25^, had any impact on *Nfil3* expression, weight gain, serum triglycerides, or gWAT (Figures 1A–1D). These findings indicate that *C. immunis* has unique functionality compared to these closely related bacterial species. We hypothesized *C. immunis* colonized mice effectively given a single oral dose of *C. immunis* had phenotypic effects that persisted for at least one week. However, while *C. immunis* was detectable in feces 12 hours after administration, it was completely absent by 48 hours (Figure 2A), a finding that suggests the bioactive principle may be present as a pre-formed component within the *C. immunis* culture used for oral treatment. Indeed, mice treated with *C. immunis* conditioned supernatant had decreased gWAT that phenocopied mice treated with *C. immunis* (Figures 2B), a finding that confirmed this metabolic effect is mediated by a molecule released by *C. immunis*. Taken together, these data indicate that the activity of *C. immunis* is dependent on a secreted molecule that is lacking in *C. symbiosum* and *C. clostridioforme*.

**Figure 2:**
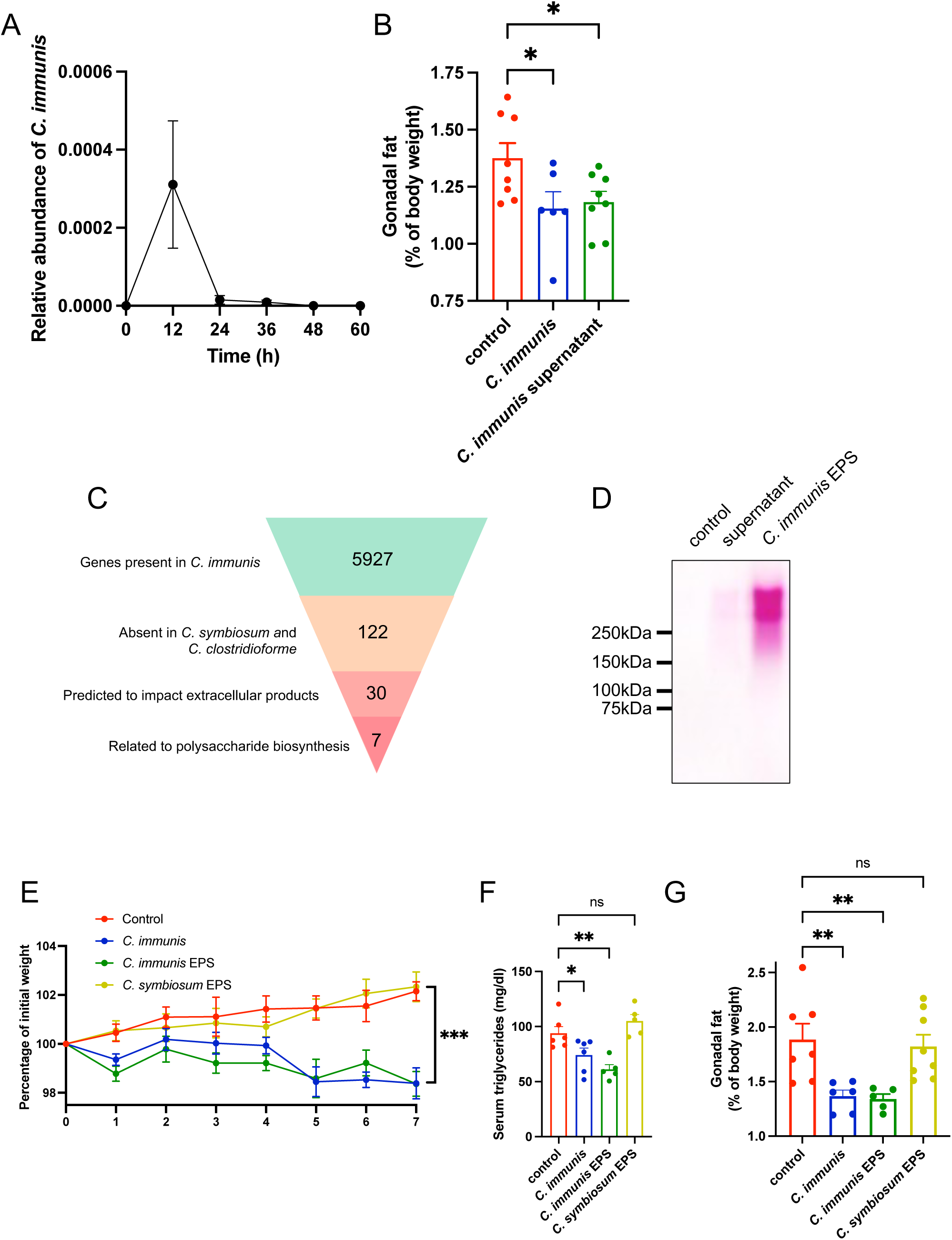
A high-molecular weight exopolysaccharide purified from *C. immunis* recapitulates its activity. (**A**) Longitudinal qPCR analysis of *C. immunis* fecal abundance in SPF C57BL/6 mice orally treated with *C. immunis.* (*n* = 10 mice). (**B**) SPF C57BL/6 mice were treated orally with sterile media (control), *C. immunis*, or conditioned supernatant from *C. immunis*. One week later, gonadal fat mass normalized to body weight was assessed. (**C**) Schematic of comparative genomic analysis for *C. immunis* genes associated with activity. (**D**) Periodic acid–Schiff-stained SDS-PAGE gel of a control, *C. immunis* supernatant, and purified *C. immunis* exopolysaccharide (EPS). The control consisted of sterile bacterial media that went through the same purification process as *C. immunis* EPS, and all samples represent an equal amount of culture volume. (**E–G**) SPF C57BL/6 mice were treated orally with sterile media (control), *C. immunis*, or EPS purified from the indicated bacteria, with body weight assessed daily for one week (**E**; *n* = 5–6 mice per group). One week after treatments, serum triglyceride (**F**) and gonadal fat mass normalized to body weight was assessed (**G**). Graphs represent means±s.e.m. \**P* < 0.05; \*\**P* < 0.01; \*\*\**P* < 0. 001; ns, not significant by one-way ANOVA with Holm-Šidák correction. For panel E, statistical comparisons were performed on the AUC. See also Figure S2

Accordingly, we performed comparative genomics to identify extracellular compounds uniquely produced by *C. immunis*. We initially identified 122 genes that were present in *C. immunis* but absent in both *C. symbiosum* and *C. clostridioforme* (Figure 2C). We manually curated this list to identify those likely to produce extracellular products, which further reduced this list to 30 genes (Figure 2C, Table S1). Intriguingly, nearly one-quarter of these genes were related to biosynthesis of an extracellular polysaccharide, a well-described mechanism by which commensal bacteria modulate host physiology^31–33^. As such, we purified from *C. immunis* supernatant a high-molecular weight exopolysaccharide (EPS; Figure 2D), which contained no appreciable protein, nucleic acid, or lipid contamination (Figures S2A–C). Critically, mice treated with this *C. immunis* EPS—but not EPS purified from *C. symbiosum*—had decreased weight gain, serum triglycerides, and gWAT compared to control animals (Figures 2E–G). These data conclusively establish that the *C. immunis* EPS is the relevant bioactive molecule that protects against metabolic disease in vivo.

### Phosphocholine modification of the *C. immunis* EPS is essential for activity

To understand the structural determinants of activity of *C. immunis* EPS, we performed glycosyl compositional analysis on EPS purified from *C. immunis* and *C. symbiosum*. The overall monosaccharide composition did not differ markedly between the two species, with both EPS structures composed primarily of mannose and glucose (Figure S3A). Although this mannan-like composition is common in plants and fungi, it is rare in bacterial polysaccharides^34,35^. Furthermore, the ^1^H-NMR spectra for both EPS samples from *C. immunis* and *C. symbiosum* were very similar with respect to the anomeric signals that form the carbohydrate portion (Figure 3A). However, the *C. immunis* EPS contained a narrow signal at 3.22 ppm suggestive of phosphocholine; this sharp singlet is absent in the *C. symbiosum* EPS, which instead has a triplet slightly downfield (Figure 3A). Of note, phosphocholine moieties on polysaccharides (e.g., capsule, lipooligosaccharides) from respiratory pathogens enhance virulence by modulating the host immune system^36^, an important precedent for an analogous role that phosphocholine may play in regulating host–commensal interactions in the gut. Upon closer examination of the genes identified as distinguishing *C. immunis* from *C. symbiosum* and *C. clostridioforme*, we found two genes that shared homology to a choline transporter (*licB*) and a phosphocholine transferase (*licC*; Table S1). These genes reside in a three-gene operon that also contains a choline kinase (*licA*; Figure S3B).

**Figure 3:**
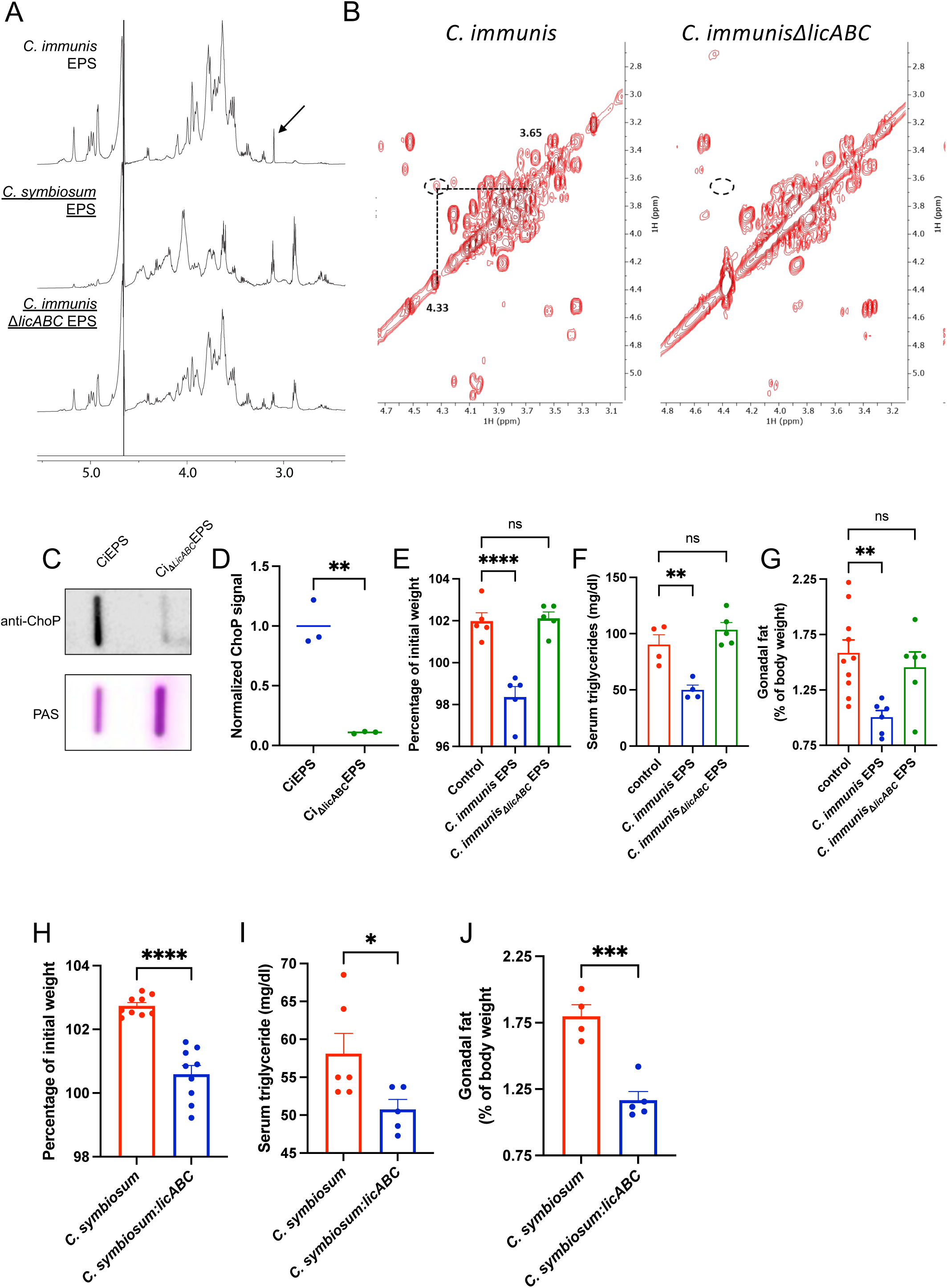
Phosphocholine modification of *C. immunis* EPS is essential for activity. **(A)** ^1^H-NMR spectra of EPS purified from *C. immunis, C. symbiosum*, or *C. immunis*Δ*licABC*. The arrow highlights a peak corresponding to phosphocholine, which is only present in EPS from *C. immunis*. **(B)** Zoomed region of 2D-COSY spectra of the *C. immunis* (left) and *C. immunis_ΔlicABC_* EPS (right) highlighting the cross peak for two of the phosphocholine protons, which is absent in the *C. immunis_ΔlicABC_* EPS. **(C)** Immunoblot of CiEPS and Ci_ΔlicABC_EPS that was probed with an antibody against phosphocholine (anti-ChoP; top blot). Samples were also visualized with periodic acid–Schiff stain (PAS; bottom blot). **(D)** Quantification of the band intensities in panel C, with values from the immunoblot normalized to that from the PAS-stained blot. (**E–G**) C57BL/6 mice were treated orally with sterile media (control), *C. immunis* EPS, or *C. immunis*_Δ*licABC*_ EPS. After one week, relative body weights (**E**), serum triglycerides (**F**), and gonadal fat mass normalized to body weight were assessed (**G**). (**H–J**) C57BL/6 mice were treated orally with *C. symbiosum* or *C. symbosum:licABC*. One week later, relative body weights (**H**), serum triglycerides (**I**), and gonadal fat mass normalized to body weight were assessed (**J**). \**P* < 0.05; \*\**P* < 0.01; \*\*\**P* < 0. 001; \*\*\*\**P* < 0. 0001; ns, not significant by unpaired Student’s t test (**D**, **H–J**) or one-way ANOVA with Holm-Šidák correction (**E–G**). See also Figure S3.

To determine whether this *licABC* operon is required for activity, we first needed to develop a system to genetically manipulate *C. immunis*, a feat that is still challenging for most commensal organisms^37^. Although CRISPR-Cas9 has been a useful approach to make isogenic mutants in some clostridial species^38,39^, expression of Cas9 was toxic to *C. immunis*, similar to its effect in many other commensal bacteria^40^. Instead, we created a suicide plasmid that contained an erythromycin resistance cassette flanked by DNA homologous to *C. immunis licA* to facilitate homologous recombination that disrupted the entire *licABC* locus (Figure S3C).

We isolated EPS from the *C. immunis*Δ*licABC* isogenic mutant (Ci_ΔlicABC_EPS) and found that its monosaccharide composition was similar to that of wild-type *C. immunis* EPS (CiEPS; Figure S3A). Heteronuclear single quantum coherence (HSQC) spectra, which shows the correlation between a carbon and hydrogen bonded directly to it, for both CiEPS and Ci_ΔlicABC_EPS confirmed that the anomeric regions were not significantly different between the two samples (Figure S3D). Combining these data with linkage analysis (Figure S3E), we were able to assign all the anomeric protons in the ^1^H-NMR (Figure S3F). However, as anticipated, ^1^H-NMR, HSQC, and correlation spectroscopy (COSY), which reveals correlations between protons that are 2–3 bonds apart, confirmed the absence of phosphocholine in Ci_ΔlicABC_EPS (Figures 3A, 3B, S3G). We further confirmed the absence of phosphocholine in Ci_ΔlicABC_EPS via an immunoblot (Figures 3C, 3D).

Thus, we used 3 orthogonal methods—structural analysis, bacterial genetics, and immune recognition—to conclusively demonstrate that *C. immunis* EPS contains phosphocholine. Consistent with phosphocholine being critical for the activity of *C. immunis* EPS, we found that mice treated with Ci_ΔlicABC_EPS had weight gain, serum triglycerides, and levels of adiposity that were indistinguishable from control animals (Figures 3E–G). These findings raise the possibility that phosphocholine—even in the absence of the polysaccharide—is sufficient to regulate VAT. However, treatment of mice with free phosphocholine had no effect on VAT (Figure S3H). To further demonstrate the critical nature of phosphocholine on EPS, we engineered *C. symbiosum* to express the *licABC* genes and thus produce a phosphocholine-modified EPS (Figure S3I). Mice treated with this bacterial gain-of-function mutant had diminished weight gain, decreased serum triglycerides, and reduced gWAT compared to the parental strain of *C. symbiosum* (Figures 3H–J). Given that neither the phosphocholine-free polysaccharide (i.e., *C. symbiosum* EPS and Ci_ΔlicABC_EPS) nor free phosphocholine can reduce VAT, our data clearly demonstrate that the phosphocholine must be attached to the EPS for its ability to modulate host metabolism.

### *C. immunis* activity requires decreased IL-22 levels

Having determined the bacterial mechanism by which *C. immunis* regulates VAT, we sought to define the host mechanisms required for its activity. Recent work established that the microbiota positively regulates *Nfil3* expression and VAT by increasing dendritic cell (DC) secretion of IL-23, which elicits more IL-22 from ILC3s^28^. In our case, *C. immunis* negatively regulates small-intestinal expression of *Nfil3* and the amount of VAT, and we hypothesized these effects are mediated through the same host-mediated pathways. However, we found that levels of IL-23 and IL-1β, the two primary cytokines secreted by DCs that modulate ILC3 function^41,42^, were not altered in *C. immunis*-treated mice (Figures S4A, S4B). In contrast, we found that *C. immunis* can directly impact ILC3 function. MNK-3 cells, an ILC3-like cell line^43^, treated with *C. immunis* EPS had decreased expression of *Il22* with no impact on expression of *Il17a* or *Csf2*, which encodes GM-CSF (Figures 4A, S4C, S4D). Notably, this activity was dependent on phosphocholine (Figure 4A).

**Figure 4:**
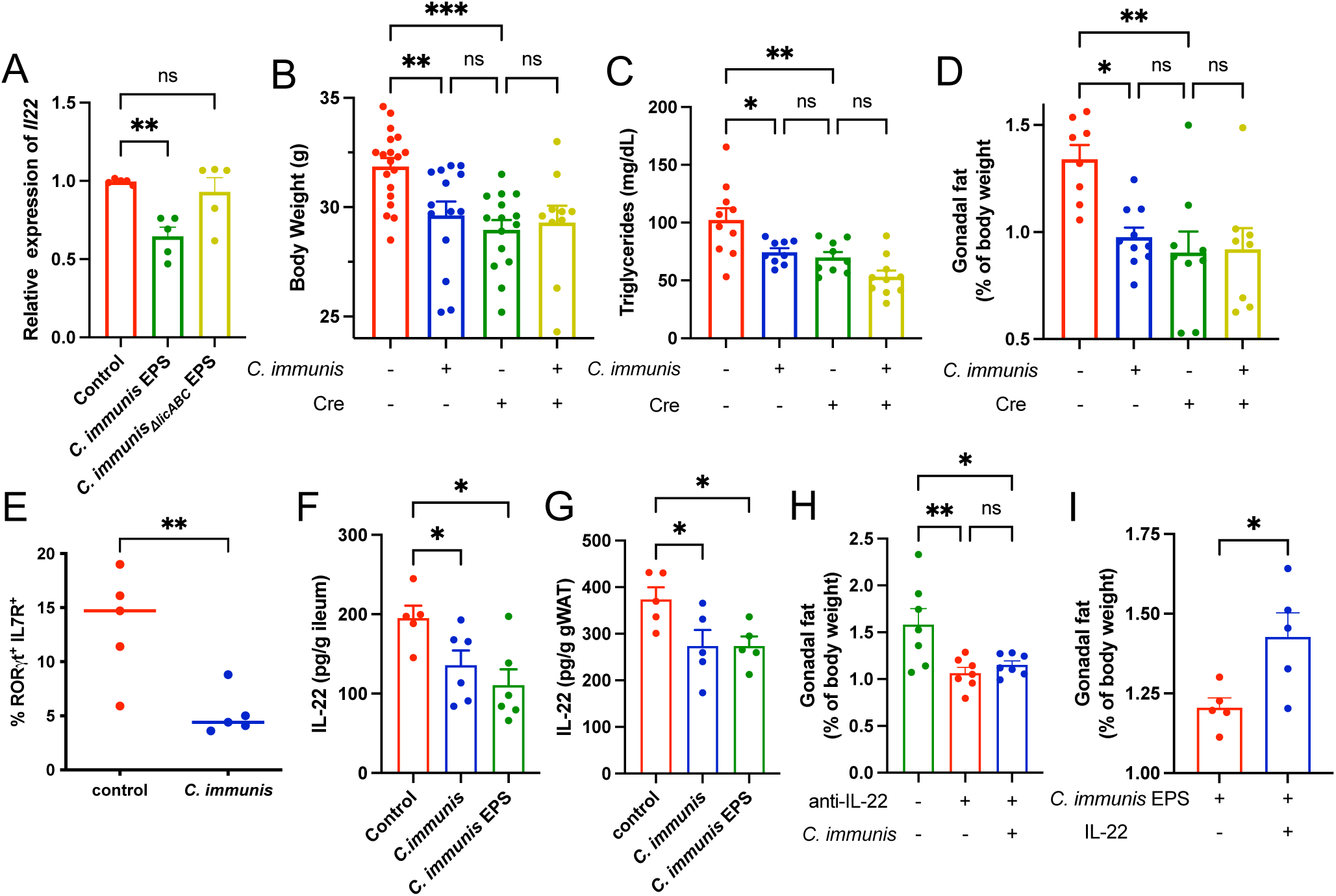
*C. immunis* activity requires decreased IL-22 levels. (**A**) qPCR analysis of *Il22* expression in MNK-3 cells treated with sterile media (control), *C. immunis* EPS, or *C. immunis*_Δ*licABC*_ EPS. **(B–D)** *Ahr^fl/fl^*(ILC3-sufficient) and *Ahr^fl/fl^; Rorc-cre* mice (ILC3-deficient) were orally treated with or without *C. immunis*, with body weights (**B**), serum triglycerides (**C**), and gonadal fat mass normalized to body weight (**D**) assessed one week later. (**E**) Flow cytometric analysis of the frequency of intestinal RORγt^+^ IL7R^+^ cells when gated on lineage negative cells (CD3^−^ CD19^−^ CD11b^−^ CD11c^−^). (**F**, **G**) IL-22 ELISA results for ileal tissue (**F**) and gWAT (**G**) from C57BL/6 mice, one week after orally treating with sterile media (control), *C. immunis*, or *C. immunis* EPS. (**H**) Gonadal fat mass normalized to body weight in C57BL/6 mice treated with or without an IL-22 neutralizing antibody one week prior, with some mice having also received *C. immunis*. (**I**) Gonadal fat mass normalized to body weight in C57BL/6 mice orally treated with *C. immunis* EPS one week prior, with one group having also received IL-22 IP daily. \**P* < 0.05; \*\**P* < 0.01; \*\*\**P* < 0.001; ns, not significant by ANOVA with Holm-Šidák correction (**A–D, F–H**) or Student’s t test (**E**, **I**). See also Figure S4.

To determine whether ILC3s are required in vivo for *C. immunis-*mediated regulation of metabolism, we generated ILC3-deficient mice by breeding *Ahr^fl/fl^*mice with *Rorc-cre* mice, as previously described^44^. These ILC3-deficient mice had lower body weights, decreased serum triglyceride levels, and reduced amounts of gWAT compared to *Ahr^fl/fl^* littermate controls (Figures 4B–D), findings that are consistent with the known role for ILC3s in obesity^28,30,45^. While *C. immunis* treatment of *Ahr^fl/fl^* mice phenocopied the effect of ILC3 deficiency, it had no effect in ILC3-deficient mice (Figures 4B–D). These findings highlight that *C. immunis* is only active when ILC3s are present, which indicates that ILC3s are required for its activity. Consistent with these findings, *C. immunis*-treated mice had reduced ILC3s in the intestine, which helps explain why these mice mirror the phenotype of ILC-deficient mice (Figures 4E, S4E). Of note, *Ahr^fl/fl^ Rorc-cre* mice also lack *Ahr* expression in T cells^44,46^. Although this defect has not been found to have significant effects in vivo^47,48^, we confirmed our results are not related to T cells by showing *C. immunis* decreases gWAT in *Rag1^−/−^* mice (Figure S4F). To further ensure there is no functional redundancy between AHR in T cells and ILC3s, we generated ILC3-deficient mice on a *Rag1^−/−^*background. We found that *Rag1^−/−^*;*Ahr^fl/fl^*;*Rorc-cre* mice treated with *C. immunis* EPS did not lose weight or gWAT compared to untreated *Rag1^−/−^*;*Ahr^fl/fl^*;*Rorc-cre* mice; in contrast, *C. immunis* EPS reduced both features in *Rag1^−/−^*;*Ahr^fl/fl^* littermate controls (Figure S4G, S4H). Taken together, these data establish that *C. immunis* regulates metabolic disease in an ILC3-dependent manner.

Based on the in vivo requirement for ILC3s and our in vitro results with MNK-3 cells, we speculated that *C. immunis* treatment decreases levels of IL-22 in vivo given that ILC3s are a major source of this cytokine^49^. Indeed, treatment with either *C. immunis* or *C. immunis* EPS significantly decreased levels of ileal and gWAT IL-22 levels without altering levels of IL-22 in iWAT (Figures 4F, 4G, S4I). Although IL-22 is critical for maintaining intestinal homeostasis^50^, *C. immunis* treatment did not adversely affect barrier integrity or ileal expression of *Reg3g* (Figures S4J, S4K), a canonical IL-22-dependent gene that helps regulate host–microbiota interactions^49,51^. To confirm this decrease in IL-22 was important for *C. immunis* activity, we used a neutralizing antibody to deplete mice of IL-22. These IL-22-depleted mice had decreased amounts of gWAT compared to control animals, and *C. immunis* lacked the ability to further reduce gWAT in these IL-22-depleted mice (Figure 4H). To complement these studies, we gave *C. immunis* EPS-treated mice exogenous IL-22, which increased small-intestinal and gWAT levels of IL-22 to normal levels (Figures S4L, S4M). Although treatment with supraphysiological levels of IL-22 has previously been associated with improved metabolic health^52^, the addition of IL-22 resulted in EPS-treated mice having higher body weights and more gWAT than mice treated only with *C. immunis* EPS (Figures 4I, S4N). In addition, treatment of *Rag1^−/−^*;*Ahr^fl/fl^*;*Rorc-cre* mice with IL-22 increased gWAT levels, establishing that the addition of IL-22 is sufficient to overcome the effect on gWAT related to ILC3 deficiency (Figure S4O). Collectively, these data demonstrate that the ability of *C. immunis* EPS to reduce VAT requires decreased numbers of ILC3s and levels of IL-22.

### *C. immunis* EPS increases metabolic activity in VAT

Although we identified *C. immunis* EPS as the bioactive molecule and defined decreased IL-22 levels as a requirement for its activity, it remains unclear how energy balance is impacted such that body weight and visceral adiposity is reduced. Accordingly, we performed indirect calorimetry studies of *Ahr^fl/fl^* mice fed a high-fat diet for 12 weeks and orally treated every week with or without *C. immunis* EPS; we also included untreated ILC3-deficient littermate mice fed a high-fat diet. We found no difference in energy intake or nutritional absorption among groups given no change in cumulative food intake, similar results with fecal bomb calorimetry, and no difference in fecal lipids (Figures 5A, S5A–C). Rather, *C. immunis* EPS-treated *Ahr^fl/fl^* mice and untreated ILC3-deficient mice had increased energy expenditure compared to untreated *Ahr^fl/fl^*mice (Figure 5B), with no change in physical activity (Figure S5D).

**Figure 5:**
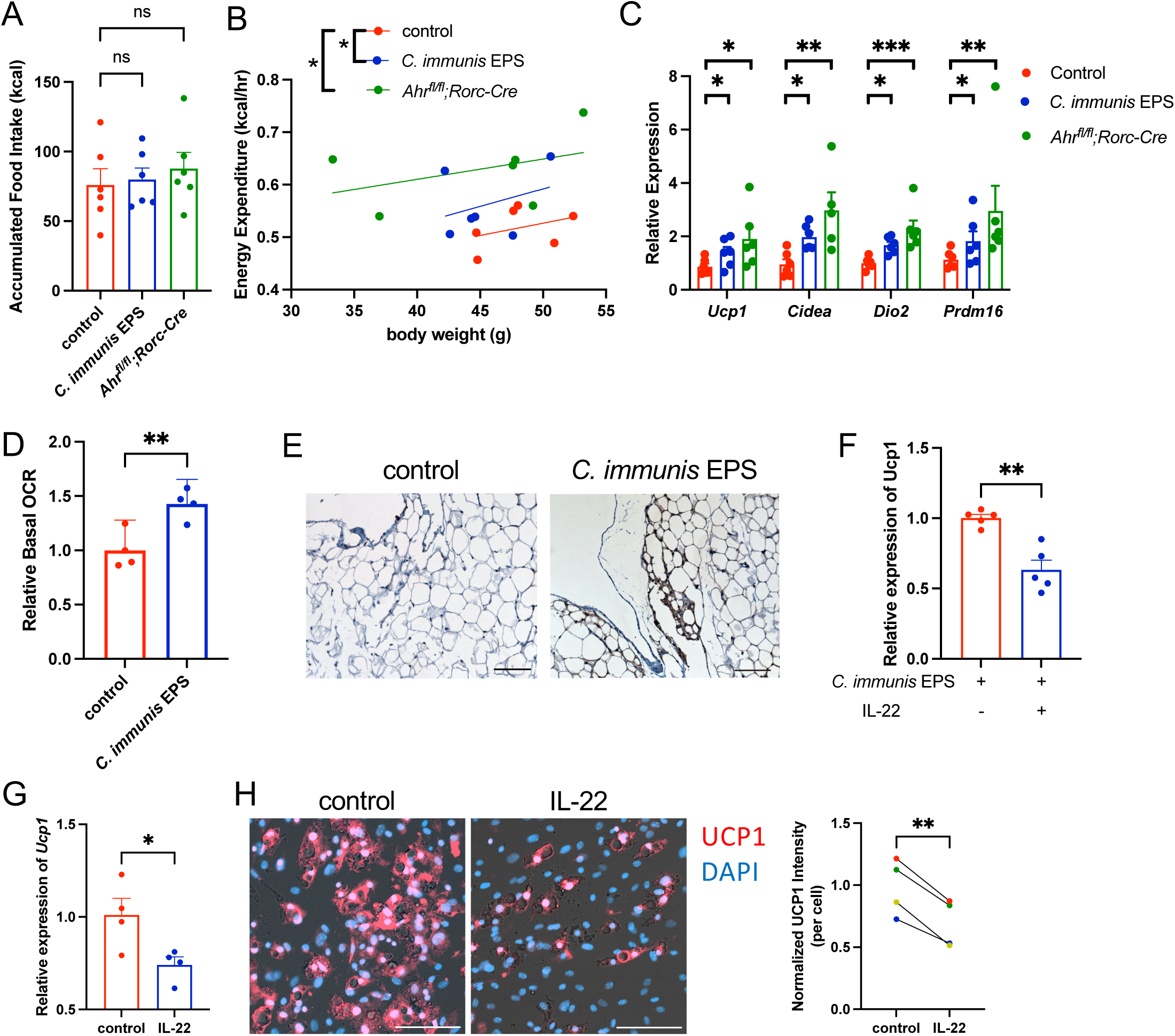
*C. immunis* EPS increases metabolic activity in VAT. (**A**, **B**) *Ahr^fl/fl^*mice orally treated with or without weekly *C. immunis* EPS and untreated *Ahr^fl/fl^; Rorc-Cre* mice were fed a high-fat diet for 12 weeks prior to indirect calorimetry studies. Accumulated food intake (**A**) and energy expenditure as a function of body weight (**B**) are shown. (**C**) qPCR analysis of the indicated genes in gWAT obtained from *Ahr^fl/fl^* mice orally treated with or without weekly *C. immunis* EPS and untreated *Ahr^fl/fl^; Rorc-Cre* mice. Mice were fed a high-fat diet for 12 weeks prior to the analysis. (**D**) Basal oxygen consumption rates (OCR) for gWAT samples normalized to the control samples. Each dot represents the average of 11–12 technical replicates. (**E**) Immunohistochemistry of UCP1 in gWAT of C57BL/6 mice treated with or without *C. immunis* EPS one week prior to sacrifice. Images are representative of findings from 3 independent mice per group. Scale bar represents 100 μm. (**F**) qPCR analysis of *Ucp1* in gWAT of C57BL/6 mice orally treated with *C. immunis* EPS one week prior, with one group having also received IL-22 IP daily. (**G–H**) The SVF obtained from gWAT was induced for beige cell differentiation in the presence or absence of IL-22, with qPCR analysis of *Ucp1* expression (**G**) and fluorescence microscopy images using an antibody against UCP1 (red) and counterstained with DAPI (blue; **H**) shown. Scale bar represents 100 μm. Each dot in the graph in panel H represents the normalized fluorescence intensity of UCP1 averaged from 5–6 fields per sample, with the color of dot and lines indicating samples that came from the same mouse. \**P* < 0.05; \*\**P* < 0.01; \*\*\**P* < 0. 001; ns, not significant by one-way ANOVA with Holm-Šidák correction (**A**, **C**), ANCOVA (**B**), Student’s t test (**D**, **F**, **G**), or paired t-test (**H**). See also Figure S5.

We speculated this decrease in VAT coupled with an increase in energy expenditure may indicate increased VAT-associated thermogenesis. Consistent with this notion, there was increased expression of several thermogenesis-related genes (*Ucp1*, *Cidea*, *Dio2*, *Prdm16*) in the gWAT of *C. immunis* EPS-treated *Ahr^fl/fl^* mice and untreated ILC3-deficient mice compared to untreated *Ahr^fl/fl^* mice (Figure 5C); these changes were not present in iWAT (Figure S5E). In line with these findings, gWAT from *C. immunis* EPS-treated mice had increased basal metabolic activity as measured by respirometry compared to control animals (Figure 5D). Immunohistochemistry revealed an increased number of UCP1^+^ multiloculated cells in gWAT isolated from *C. immunis* EPS-treated mice compared to control animals (Figure 5E). Moreover, giving exogenous IL-22 to *C. immunis* EPS-treated mice decreased *Ucp1* expression in gWAT (Figure 5F), a finding that demonstrates IL-22 negatively regulates expression of *Ucp1* in gWAT. Taken together, these findings suggest that *C. immunis* EPS increases energy expenditure by recruiting UCP1^+^ cells in VAT and increasing its metabolic activity.

To determine whether IL-22 directly influences the induction of UCP1, the stromal vasculature fraction (SVF) of iWAT and gWAT was induced for beige cell differentiation in the presence or absence of IL-22. Consistent with previous work^53^, the addition of IL-22 increased the expression and protein levels of UCP1 in iWAT SVF (Figures S5F, S5G). Although VAT is generally more resistant to recruitment of UCP1^+^ cells than other WAT depots^54–56^, SVF isolated from gWAT was still able to be induced into UCP1^+^ cells (Figure 5F), albeit less robustly than SVF from iWAT. In stark contrast to its role in stimulating UCP1 levels in iWAT SVF, the addition of IL-22 reduced expression and protein levels of UCP1 in differentiated gWAT SVF (Figure 5G, 5H). In line with these findings, administration of IL-22 to otherwise untreated healthy mice resulted in increased *Ucp1* expression in iWAT but not gWAT (Figure S5H). Thus, the reduction in gWAT IL-22 levels driven by *C. immunis* EPS results in the increased recruitment of UCP1^+^ cells in gWAT, which helps explain the increased energy expenditure, reduction in body weight, and decreased visceral adiposity.

### The abundance of phosphocholine biosynthesis genes is lower in humans with metabolic disease

Given that modification of secreted molecules with phosphocholine is a common strategy by which bacteria in the respiratory tract evade detection and clearance by the host immune system, we postulated that this strategy may also be used by other intestinal commensal bacteria. Using the *C. immunis licABC* gene sequence as a reference, we identified numerous taxonomically diverse gut commensal bacteria that contain a *licABC* locus (Figure 6A). Intriguingly, the gene sequences for bacteria primarily found in the respiratory tract cluster separately from intestinal organisms, which itself forms two distinct clades. Given the phylogeny of the *licABC* gene sequences from the gut commensal bacteria is distinct from the bacterial taxonomic relationships, it is possible the *licABC* locus is spread via horizontal gene transfer. Along these lines, it is interesting to note that the *C. immunis licABC* operon is flanked by mobile genetic elements (Figure S3B), which may help explain why closely related taxa lacked these genes.

**Figure 6:**
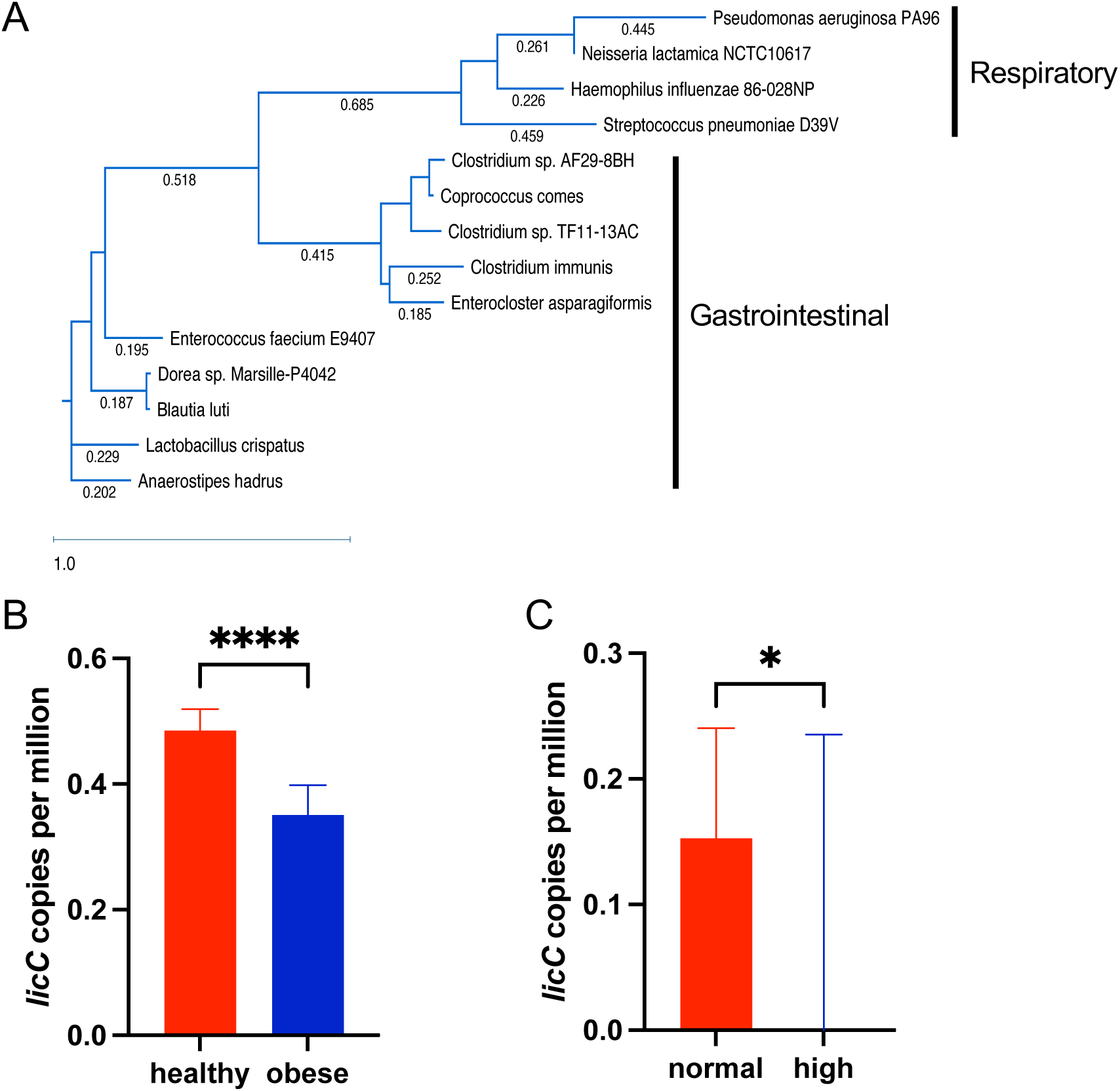
The abundance of phosphocholine biosynthesis genes is lower in humans with metabolic disease. **(A)** A randomized axelerated maximum likelihood (RAxML) phylogenetic tree depicting *licABC* genes identified in *C. immunis* and other gut commensal bacteria is shown, with the *licABC* genes from four bacteria primarily found in the respiratory tract added in for reference. **(B)** The abundance of *licC* in human fecal metagenomic data is shown for individuals with a healthy (< 25 kg/m^2^; *n* = 2021) or obese BMI (≥ 30 kg/m^2^; *n* = 816). **(C)** The abundance of *licC* in human fecal metagenomic data is shown for individuals with normal (<150 mg/dL; *n* = 193) or high serum levels of triglycerides (≥200 mg/dL; *n* = 22). Data are represented as medians with the 95% confidence interval. For panel C, the median *licC* abundance for individuals with hypertriglyceridemia is 0. \**P* < 0.05; \*\*\*\**P* < 0.0001 by Mann-Whitney test (**B, C**). See also Figure S6.

To determine whether our findings are generalizable to metabolic disease in humans, we analyzed publicly available shotgun metagenomic data for the abundance of these phosphocholine biosynthesis genes. We focused on *licC*, which generates the critical CTP-phosphocholine substrate required for decoration of bacterial polysaccharides, as this gene is least commonly found in gut commensal bacteria^57^. Analyzing data from 4,145 otherwise healthy individuals (pooled from 11 studies that included individuals from 9 different countries) for whom BMI data was available^58^, we found an overall negative correlation between the abundance of *licC* and BMI (Figure S6A). More specifically, individuals with a normal BMI (< 25 kg/m^2^) had a higher abundance of *licC* genes than obese individuals (BMI ≥ 30 kg/m^2^; Figure 6B). Additionally, using a different collection of 247 individuals (pooled from 4 studies that included individuals from 6 different countries) for whom serum triglyceride data was available^58^, we similarly found an overall negative correlation between the abundance of *licC* and serum triglyceride levels (Figure S6B), with those with normal triglyceride levels (<150 mg/dL) having a higher abundance of *licC* genes than individuals with hypertriglyceridemia (≥200 mg/dL; Figure 6C). Taken together, these results strongly suggest that our murine findings of a *licABC*-modified bacterial compound regulating triglycerides, weight, and VAT are generalizable to humans.

## DISCUSSION

Obesity is a deepening global health crisis, with ∼4 million deaths worldwide annually attributed to individuals being overweight or obese^2^. Given increasing recognition that the microbiota is a critical regulator of lipid metabolism and body composition, microbiome-derived therapeutics may provide scalable and cost-effective solutions to metabolic disease. Here we identify and characterize a specific human commensal bacterium-derived EPS that protects against obesity by recruiting UCP1^+^ cells to VAT and increasing overall energy expenditure.

Although comparisons of germ-free and conventionalized animals revealed the net effect of the microbiota is to promote the expansion of VAT^22^, treatment with *C. immunis*—similar to a few other commensal bacteria^20,59–61^—results in a decreased amount of VAT. This finding highlights the complexity of microbiome-mediated modulation of metabolic conditions where different organisms have completely opposite effects, with the resulting physiologic changes representing an emergent phenotype. Although the specific bacterial mechanism(s) that leads to altered adiposity has not previously been characterized, there are limited examples suggesting bacterial viability and colonization are required for decreasing adiposity^20,60^. In contrast, *C. immunis*, which does not colonize SPF animals when administered as a probiotic, does so by secreting an immunomodulatory EPS into its environment.

Given that bacteria commonly synthesize high-molecular weight polysaccharides that are either released or secreted into its milieu^62^, it is not clear what drives specificity in the host response to them. Assuming only 1% of commensal bacteria produce host-accessible polysaccharides, an estimated average molecular weight of 100 kDa for the polysaccharides, and—based on our findings with *C. immunis*—that there is ∼0.1 g polysaccharide for every 10^10^ cfu bacteria, there are ∼10^9^ bacterial polysaccharides per intestinal epithelial cell given current estimates for the number of commensal bacteria and host cells^63^. Even if each of these assumptions were 10–100 fold too high, there would still be an overwhelming number of bacterial polysaccharides per epithelial cell, which begs the question of how the host distinguishes between all these molecules and what—from the bacterial side—drives specificity.

Indeed, polysaccharides are the most chemically diverse molecule: there are >20 “common” constituent sugars (and many more uncommon sugars), variable linkage patterns (α or β) that can occur at different positions on each sugar substituent, and modifications that include other common biological molecules (e.g., lipids, proteins)^64^. Most of this structural diversity remains unexplored. Bacteria are thought to contain the most diverse (and mostly unexplored) polysaccharides^65,66^, owing, in part, to the sheer number of different bacterial species, the inclusion of more uncommon sugars, and a diverse array of modifications. Given that these bacterial glycans are often present on the outermost surface, they play critical roles in mediating host–bacterium interactions, where they facilitate colonization, invasion, and modulate the adaptive and innate arms of the immune system^33,66^.

We found that *C. immunis* and *C. symbiosum* produced structurally similar, though functionally divergent, EPS molecules that differed primarily in the presence of a phosphocholine moiety. Polysaccharides modified with phosphocholine are present in taxonomically diverse microbes that colonize humans, particularly bacteria that are present in the respiratory tract^36,67,68^. Phosphocholine-containing bacterial polysaccharides contribute to bacterial colonization^69–71^, persistence^72,73^, and virulence^72,74^. Moreover, they can facilitate both evasion from, and recognition by, the immune system^36,75,76^. Genes in the *lic* operon are responsible for adding phosphocholine to polysaccharides; however, neither the *licABC* locus nor phosphocholine-containing polysaccharides were previously found in clostridial species^36,57^. To better define the role of the phosphocholine moiety in the *C. immunis* EPS, we developed means to genetically manipulate *C. immunis* and inactivated the *licABC* locus. We found that this isogenic mutant, which produces an EPS identical to the parent isolate but lacking phosphocholine, failed to modulate immune responses or visceral adiposity. In addition, we engineered *C. symbiosum* to express *licABC* and produce a phosphocholine-modified EPS that was now able to reduce visceral adiposity and weight gain. Taken together, these findings conclusively demonstrate the critical role for the presence of phosphocholine on EPS in modulating metabolic disease.

Our identification of *licABC* as the bacterial genes required for activity of the *C. immunis* EPS enabled us to identify this operon in several other commensal gut bacteria. These gut bacterial *licABC* genes may have eluded prior identification given their sequences are quite distinct from that of respiratory pathogens, in which *licABC* genes have been extensively characterized^36^. It is likely that these other gut commensal bacteria decorate some surface-associated molecule with phosphocholine, similar to what has been observed with *licABC*-containing respiratory pathogens^36^. It is possible that modification of common commensal-associated molecular patterns with small haptens, such as phosphocholine, may help create a “combinatorial code” that provides increased specificity to host–microbiota interactions and thus fine-tunes physiological perturbations.

It is interesting to speculate that the importance of this phosphocholine moiety is related to the fact that it contains both positively (i.e., choline) and negatively charged (i.e., phosphate) groups. Bacterial polysaccharides are generally neutral or negatively charged, with few containing a positive charge^33,77^. However, bacterial polysaccharides that contain a zwitterionic motif like the *C. immunis* EPS are known to modulate the immune system^77–80^. *Bacteroides fragilis* polysaccharide A is the archetypal example of an immunomodulatory zwitterionic polysaccharide (ZPS), where chemical neutralization of either charged group results in a functionally inactive molecule^77^. These previously described ZPSs are presented via the MHCII pathway such that they activate CD4^+^ T cells^80,81^. In contrast, our in vitro findings suggest that the *C. immunis* EPS may directly modulate the function of ILC3s in a manner dependent on its zwitterionic phosphocholine moiety.

Specifically, *C. immunis* EPS-treated mice have decreased small-intestinal and VAT levels of IL-22, with no change detected in subcutaneous adipose tissue (SAT). This decrease in IL-22 levels is required for *C. immunis* EPS-mediated protection against metabolic disease, which we demonstrated by neutralizing IL-22 and, alternatively, by giving exogenous IL-22 to *C. immunis* EPS-treated animals. Although IL-22 is generally regarded as a key cytokine in regulating VAT, there is ongoing controversy regarding its specific role. Initial studies found IL-22 protects against metabolic disease^52,82,83^, but others have found that it instead promotes metabolic dysfunction^28,30,84^. In contrast, *Il22^−/−^* mice had no metabolic phenotypes^52,85^, a finding that suggests IL-22 may not be essential to metabolic health. Our results help resolve these enduring discrepancies. Since we found that SAT and VAT have disparate responses to IL-22, the differences observed in these earlier studies may relate, in part, to which tissues were more impacted by the changes in IL-22 levels.

Given that UCP1^+^ adipocytes help burn energy and protect against metabolic diseases, there has been a longstanding interest in therapies that can induce these cells in VAT^86^. However, the canonical signals that induce these cells (e.g., exposure to cold temperature, β-adrenergic agonists, other hormone-like compounds) are far more effective in SAT^87–89^, with VAT being relatively resistant to these agents^54–56^. We found that the *C. immunis* EPS and the resulting decreased local levels of IL-22 drives recruitment of UCP1^+^ cells specifically in VAT without impacting SAT, thus providing new insights into the maintenance and remodeling of VAT. The reason for the difference in biological responses between SAT and VAT remains unknown, though we speculate it may relate to VAT being covered by a layer of mesothelial cells. Interestingly, these VAT-associated mesothelial cells in both humans and mice express the IL-22 receptor and may therefore influence how IL-22 affects VAT remodeling^90^. This increase in UCP1^+^ adipocytes is associated with increased metabolic activity present in gWAT from *C. immunis* EPS-treated mice. Although we do not know how much of this increase in metabolic activity is dependent on UCP1 versus other mechanisms (e.g., futile cycling of creatine, calcium, and/or triglycerides), a visceral fat specific deletion of *Zfp423*, which serves as a transcriptional “brake” on the adipocyte thermogenic program and affects UCP1-dependent and independent pathways, shows many of the same phenotypes as *C. immunis* EPS-treated mice^91^. Future work will characterize more fully the specific metabolic pathways that contribute to *C. immunis* EPS-mediated decreases in VAT.

Considered together, our work couples biochemical and genetic approaches to characterize a commensal bacterium-derived, phosphocholine-containing EPS that protects against obesity in an IL-22-dependent manner. Our findings offer the possibility for a microbiome-derived product that treats obesity and related comorbidities. More broadly, our work highlights that as the ability to culture and genetically manipulate commensal organisms continues to advance, demonstrating the genetic, structural, and mechanistic basis for how the microbiome impacts host physiology will become more commonplace, just as it has become de rigueur in microbial pathogenesis.

### Limitations of the study

We have found that the effects of the *C. immunis* EPS persist for at least a week after oral treatment; however, it remains unclear whether the EPS itself remains present for this entire duration or if the immunological effects remain durable even after the EPS is lost. Future structure–function studies are needed to determine whether the polysaccharide sequence—in addition to the presence of phosphocholine—is also critical for activity. Although our data demonstrating the same polysaccharide with or without a phosphocholine modification differ in functionality may suggest otherwise, it is formally possible that the *C. immunis* EPS is modulating the microbiome in a way that is ultimately causing the improvements in metabolic health. Furthermore, although our in vitro studies suggest that *C. immunis* EPS can modulate ILC3 activity directly, we have not ruled out the possibility that some other cell type plays an important role in recognizing the EPS upstream of ILC3s. Finally, although the abundance of the *licC* gene is higher in humans without metabolic disease, it is not yet clear whether this difference results in the same mechanistic pathway as we uncovered in mice.

## ACKNOWLEDGEMENTS

We thank the Duke Gnotobiotic Core, the DLAR Breeding Core, and the Sequencing and Genomic Technologies Core for technical assistance; Dustin R. Middleton for advice on polysaccharide enrichment; Parastoo Azadi and Christian Heiss at the Complex Carbohydrate Research Center (University of Georgia), which is supported by the US Department of Energy under award DE-SC0015662, for performing structural analyses of purified exopolysaccharides; the University of Michigan Animal Phenotyping Core, which is supported by NIH grants 1U2CDK110768, DK020572, and DK089503, for performing fecal bomb calorimetry; and the Nutrition Obesity Research Center at the University of North Carolina Chapel Hill, which is supported by NIH grant P30DK056350, for performing the respirometry experiments. CYT received support from American Heart Association pre-doctoral fellowship AHA 897275, a National University of Singapore Development Grant, a Tan Kah Kee Postgraduate Scholarship, and a Triangle Center for Evolutionary Medicine Graduate Student Award. NKS received support from NIH awards R03 AI142341, R56 DK140174, and P30 DK034987, the Hartwell Foundation Individual Biomedical Research Award, the Gilhuly Accelerator Fund, a Development Award from the Duke Microbiome Center, and as a Duke Translating Duke Health Scholar and a Whitehead Scholar.

## AUTHOR CONTRIBUTIONS

Conceptualization: CYT, NKS

Methodology: CYT, YL, DJ, BST, NKS

Investigation: CYT, YL, MVR

Formal analysis: DJ

Visualization: CYT, NKS

Funding acquisition: CYT, NKS

Project administration: CYT, NKS

Supervision: NKS

Writing – original draft: CYT, NKS

Writing – review & editing: CYT, YL, DJ, BST, MVR, NKS

## SUPPLEMENTAL INFORMATION

Document S1. Figures S1–S6 and Table S1

## SUPPLEMENTARY FIGURE LEGENDS

**Figure S1. *C. immunis* inhibits intestinal expression of lipid metabolism genes without affecting lean mass, related to** Figure 1. (**A**) Volcano plot depicting differentially expressed genes in colonic tissue of C57BL/6 MMb mice treated orally with or without *C. immunis* (*n* = 3 mice per group). Labeled genes are related to lipid uptake and metabolism. (**B–C**) qPCR analysis of jejunal *Cd36* (**B**) and *Scd1* expression (**C**) from C57BL/6 MMb mice treated orally with or without *C. immunis*. (**D**) Immunofluorescence images of NFIL3 (red) in small-intestinal tissue obtained from SPF C57BL/6 mice treated with or without *C. immunis*. Samples are also stained with DAPI (blue). NFIL3 fluorescence intensity (in arbitrary units) was quantified in 3–5 fields from each of 4 independent samples. (**E**) Gene set enrichment analysis for NFIL3-dependent genes in the RNAseq dataset depicted in panel A. NES, normalized enrichment score. (**F**) AUC for the body weights displayed in Figure 1B. (**G**) Absolute values of body weight for normalized data displayed in Figure 1B. (**H**) Absolute values of gWAT mass for normalized data displayed in Figure 1D. (**I**) Lean mass detected by DEXA scans in mice treated with or without *C. immunis*. (**J**) AUC for the serum glucose levels displayed in Figure 1H. Graphs represent means±s.e.m. \**P* < 0.05; \*\**P* < 0.01; \*\*\**P* < 0. 001; ns, not significant by unpaired Student’s t test (**B–D, I–J**) or one-way ANOVA with Holm-Šidák correction (**F**, **H**).

**Figure S2. Purified *C. immunis* EPS is devoid of significant protein, nucleic acid, or lipid contamination, related to** Figure 2. (**A**) Silver-stained SDS-PAGE of a control, *C. immunis* supernatant, and *C. immunis* EPS. The control included sterile bacterial media that went through the same purification process as *C. immunis* EPS, and all samples represent an equal amount of culture volume. (**B**) Safe DNA-stained agarose gel of *C. immunis* EPS. Two different amounts of the ladder were included to demonstrate sensitivity. (**C**) Sudan Black B-stained PAGE gel of *E. coli* lipopolysaccharide and *C. immunis* EPS.

**Figure S3. The polysaccharide components of the CiEPS and Ci_ΔlicABC_EPS are largely similar, related to** Figure 3. (**A**) Glycosyl composition analysis of EPS isolated from *C. immunis*, *C. symbiosum*, and *C. immunis* Δ*licABC*. (**B**) A schematic of the *C. immunis* genetic locus containing *licABC* and flanking genes. The predicted promoter is indicated by the arrow. (**C**) A schematic of the genetic strategy used to delete *licABC* in *C. immunis*. *E. coli* was used to conjugally transfer into *C. immunis* a suicide plasmid containing an erythromycin resistance gene (*ErmB*) flanked by homology arms to *licA*. Erythromycin-resistant (Ery^R^) *C. immunis* mutants contained the *ErmB* gene and an 80-bp deletion in *licA*. (**D**) Anomeric regions in the HSQC spectra of CiEPS (left) and Ci_ΔlicABC_EPS (right). The anomeric CH are labelled with letters in the *C. immunis* EPS spectra. (**E**) Linkage analysis of EPS isolated from *C. immunis* and *C. immunis* Δ*licABC*. (**F**) Zoomed anomeric region in the ^1^H NMR spectra of *C. immunis* (top) and *C. immunis_ΔlicABC_* EPS (bottom), with tentative assignments of the anomeric protons (as labeled in panel D). Superscript a and b denote residues within a linear or branched region, respectively. (**G**) HSQC spectra of *C. immunis* EPS (left) and *C. immunis_ΔlicABC_* EPS (right) that highlight the C–H interactions from the phosphocholine moiety. The dashed circles in the Ci_ΔlicABC_EPS spectrum reveals where these interactions should appear if the phosphocholine were present. (**H**) C57BL/6 mice were orally administered water (control) or phosphocholine (ChoP), with the gonadal fat mass normalized to body weight assessed one week later. (**I**) Immunoblot of EPS purified from *C. symbiosum* and *C. symbiosum:licABC* that was probed with an antibody against phosphocholine. ns, not significant by Student’s t test.

**Figure S4. The *C. immunis* EPS decreases IL-22 specifically in the small intestine and VAT without major adverse effects, related to Figure 4. (A, B) ELISA analysis of IL-23** (**A**) and IL-1β (**B**) in intestinal tissue of mice orally treated with or without *C. immunis*. (**C–D**) qPCR analysis of *Csf2* (**C**) and *Il17* (**D**) in MNK-3 cells treated with sterile media (control*), C. immunis* EPS, or *C. immunis*Δ*licABC*. (**E**) Flow cytometric analysis of the absolute number of intestinal RORγt^+^ IL7R^+^ cells when gated on lineage negative cells (CD3^−^ CD19^−^ CD11b^−^ CD11c^−^). (**F**) *Rag1^−/−^* mice were orally treated with or without *C. immunis*, and gonadal fat mass normalized to body weight was assessed one week later. (**G–H**) *Rag1^−/−^; Ahr^fl/fl^*and *Rag1^−/−^; Ahr^fl/fl^; Rorc-Cre* littermate mice were treated orally with or without *C. immunis* EPS, with daily weight change shown (**G**). One week later, gonadal fat mass normalized to body weight was assessed (**H**). (**I**) ELISA analysis of IL-22 in iWAT from C57BL/6 mice, one week after orally treating with sterile media (control), *C. immunis*, or *C. immunis* EPS. (**J**) The concentration of FITC-dextran in the serum of C57BL/6 mice 4 hours after oral gavage of FITC-dextran. Some mice had been treated orally with *C. immunis* one week prior. (**K**) qPCR analysis of ileal *Reg3g* expression in mice that had been treated orally with or without *C. immunis* one week prior. (**L–N**) Mice were treated orally with *C. immunis* EPS, with some mice having also received IL-22 IP daily. One week after treatment with *C. immunis* EPS, IL-22 levels were measured in the ileum (**L**) and gWAT (**M**), and relative body weights were assessed (**N**). (**O**) *Rag1^−/−^; Ahr^fl/fl^; Rorc-Cre* mice were treated with or without daily IL-22 IP injections, with gonadal fat mass normalizd to body weight assessed after one week. \**P* < 0.05; \*\**P* < 0.01; ns, not significant by Student’s t-test (**A**, **B**, **E**, **F**, **J–O**) or one-way ANOVA with Holm-Šidák correction (**C, D**, **G–I**). For panel G, statistical comparisons were performed on the AUC.

**Figure S5. *C. immunis* EPS does not affect intestinal nutrient absorption or expression of thermogenesis-related genes in iWAT, related to Figure 5**. (**A–B**) *Ahr^fl/fl^* mice orally treated with or without weekly *C. immunis* EPS and untreated *Ahr^fl/fl^; Rorc-Cre* mice were fed a high-fat diet for 12 weeks, after which stool samples were analyzed for fecal energy (**A**) and lipids (**B**). (**C**) Food intake was measured for 6 days in *Ahr^fl/fl^*mice orally treated with or without weekly *C. immunis* EPS and untreated *Ahr^fl/fl^; Rorc-Cre* mice fed a normal chow diet. (**D**) Ambulatory activity was determined in metabolic cage studies. (**E**) qPCR analysis of the indicated thermogenesis-related genes in iWAT obtained from C57BL/6 mice treated with or without *C. immunis* EPS. (**F**–**G**) The SVF obtained from iWAT was induced for beige cell differentiation in the presence or absence of IL-22, with qPCR analysis of *Ucp1* expression (**F**) and fluorescence microscopy images using an antibody against UCP1 (red) and counterstained with DAPI (blue; **G**) shown. Scale bar represents 100 μm. Each dot in the graph in panel G represents the normalized fluorescence intensity of UCP1 averaged from 5–6 fields per sample, with the color of dot and lines indicating samples that came from the same mouse. (**H**) qPCR analysis of *Ucp1* in gWAT and iWAT of C57BL/6 mice treated with or without IL-22 IP daily for 7 days. \**P* < 0.05; \*\**P* < 0.01; ns, not significant by one-way ANOVA with Holm-Šidák correction (**A–D**), Student’s t-test (**E–F, H**), or paired t-test (**G**).

**Figure S6. The abundance of the *licC* gene in fecal metagenomic data is negatively correlated with metabolic disease in humans, related to Figure 6**. (**A**) The abundance of *licC* gene copies is plotted against the individual’s BMI (*n* = 4,145). (**B**) The abundance of *licC* gene copies is plotted against the individual’s serum triglyceride level (*n* = 247). The Spearman’s ρ and *p* values are shown for each panel.

## STAR METHODS

### Mice

C57BL/6J (stock #000664), *Ahr^fl/fl^* (stock #006203)^92^, *Rorc-cre* (stock #022791)^46^, and *Rag1^−/−^* mice (stock #002216)^93^ were initially obtained from The Jackson Laboratory. All animals were bred and maintained in the animal facility at Duke University. Gnotobiotic MMb mice were originally obtained from Dennis Kasper (Harvard University)^94^; these mice were bred and maintained as C57BL/6 MMb mice in vinyl isolators in the Duke Gnotobiotic Core. Experimental manipulation of gnotobiotic mice was performed in autoclaved, individually ventilated cages in which a.nimals received autoclaved food (LabDiet 5K67) and water. In some experiments, mice were fed a high-fat diet (TestDiet Western Diet AIN-76A; 4.49 kcal/g; 40% of calories from fat). Mice used in experiments were sex- and age-matched and drawn randomly from the same litter, when feasible. All procedures were approved by Duke’s Institutional Animal Care and Use Committee and were conducted in accordance with National Institutes of Health guidelines.

### Bacteria

*C. immunis* was obtained from Dennis Kasper (Harvard University)^25^. *C. immunis*, *C. symbiosum* (DSM 29356), and *C. clostridioforme* (DSM 933) were grown in peptone yeast glucose (PYG) broth (Anaerobe Systems) or on brain heart infusion-supplemented (BHI-S) agar plates (ATCC Medium 1293). All clostridial strains were grown in an anaerobic chamber (Coy laboratories) with 2.5% H_2_ and 0 ppm O_2_ at 37 °C. *Escherichia coli* S17.Pir was grown aerobically in LB at 37°C.

### RNA-sequencing and analysis

One week after orally administering *C. immunis* (200 μl containing ∼10^8^ colony-forming units [CFU]) or, as a control, sterile PYG media (200 μl) to male C57BL/6 MMb mice (12 weeks of age), the proximal 2 cm of colon was collected, frozen immediately in liquid nitrogen, and stored at –80 °C until needed. Tissues were homogenized in Trizol (Invitrogen) using a bead-beating approach, and RNA was purified according to the manufacturer’s instructions. RNA Integrity Number (RIN) scores for all samples used for RNA-seq were >7.0 as assessed on a TapeStation 2200 (Agilent). cDNA libraries were generated using a mRNA HyperPrep kit (KAPA) and sequenced on a NovaSeq6000 (Illumina; S4 flow cell with 150bp paired-end reads).

FastQC (version 0.11.9) was used to assess read quality and to perform trimming, STAR (release 2.7.9a) was used to align reads to the mouse genome mm10 (GRCm39)^95^, and FeatureCounts was used to generate a count table^96^. Differential gene analysis was performed with DESeq2 (version 1.28.1)^97^, and a volcano plot was generated with EnhancedVolcano. A preranked gene set enrichment analysis was performed using GSEA 4.3.2 using a previously described NFIL3-dependent gene set as input^28,98^. R (version 4.0.0) was used for all analyses.

### qPCR analysis

RNA was isolated from tissue as described above. For cells, Trizol was added to samples, and RNA was purified according to the manufacturer’s instructions. cDNA was prepared using random hexamers and the High-Capacity cDNA synthesis kit (Applied Biosystems), and qPCR was performed on a Step One Real Time PCR system (Applied Biosystems) with iTaq Universal SYBR Green Supermix (Bio-Rad). The following primers were used: *Gadph* fwd: ACCACAGTCCATGCCATCAC; *Gadph* rev: TCCACCACCCTGTTGCTGTA; *Nfil3* fwd: CTTTCAGGACTACCAGACATCCAA; *Nfil3* rev: GATGCAACTTCCGGCTACCA; *Cd36* fwd: TCATATTGTGCTTGCAAATCCAA; *Cd36* rev: TGTAGATCGGCTTTACCAAAGATG; *Scd1*fwd: CTTCTTCTCTCACGTGGGTTG; *Scd1* rev: CGGGCTTGTAGTACCTCCTC; *Il23* fwd: CATGCTAGCCTGGAACGCACAT; *Il23* rev: ACTGGCTGTTGTCCTTGAGTCC; *Il22*fwd: GCTTGAGGTGTCCAACTTCCAG; *Il22* rev: ACTCCTCGGAACAGTTTCTCCC; *Csf2* fwd: AACCTCCTGGATGACATGCCTG; *Csf2* rev: AAATTGCCCCGTAGACCCTGCT; *Il17a*fwd: ATCCCTCAAAGCTCAGCGTGTC; *Il17a* rev: GGGTCTTCATTGCGGTGGAGAG; *Reg3g* fwd: CCTTCCTCTTCCTCAGGCAAT; *Reg3g* rev: TAATTCTCTCTCCACTTCAGAAATCCT; mUCP1 fwd: CAAAAACAGAAGGATTGCCGAAA; mUCP1 rev: TCTTGGACTGAGTCGTAGAGG; mPrdm16 fwd: CCCCACATTCCGCTGTGAT; mPrdm16 rev: CTCGCAATCCTTGCACTCA; mCidea fwd: TGCTCTTCTGTATCGCCCAGT; mCidea rev: GCCGTGTTAAGGAATCTGCTG; mDio2 fwd: CAGTGTGGTGCACGTCTCCAATC; mDio2 rev: TGAACCAAAGTTGACCACCAG. The comparative Ct method was used to quantify transcripts that were normalized with respect to *Gapdh*. For assessment of *C. immunis* persistence in mice, fecal DNA was isolated as previously described^25^, and qPCR was performed using the following primers: *C. immunis* fwd: TACAACTACTCTACCACATTGACTT; *C. immunis* rev: GCGGCGGGTTTAATTTCAGA; Universal fwd: CCTACGGGAGGCAGCAG; Universal rev: ATTACCGCGGCTGCTGGCA. The comparative Ct method was used to quantify transcripts that were normalized with respect to the universal primer results.

### Assessment of adiposity, serum chemistries, and cytokine levels

Mice (13–18 weeks of age) were orally administered either ∼10^8^ CFU of bacteria, 200 μl of conditioned bacterial supernatant, 4 mg of exopolysaccharide, or sterile PYG media. One week later, serum, and various fat depots (gonadal, retroperitoneal, inguinal, and/or interscapular brown fat) were collected. For gonadal fat, the fat pads surrounding the ovaries and fallopian tubes were collected in female mice, while the epididymal fat pads were collected in male mice. The weight of these fat depots were normalized to the animal’s body weight at time of sacrifice. Serum triglycerides were assessed either in the Hormone and Conventional Analyte Lab in the Duke Molecular Physiology Institute or by using a triglyceride colorimetric assay kit (Cayman). Dual energy X-ray absorptiometry (DEXA) experiments were performed in the Biomedical Research Imaging Center at the University of North Carolina, Chapel Hill using a Lunar PIXImus (GE Lunar Corp.). In experiments assessing the impact of *C. immunis* in the context of a high-fat diet, mice were orally administered either ∼10^8^ CFU of bacteria or 0.25 mg of EPS each week, and serum and fat depots were collected after 12 weeks. Glucose tolerance was assessed by administering glucose (2 g/kg body weight IP) to mice that had been fasted for 6 hours, and blood glucose values were measured at different times using an Accu-Chek (Roche). For measurement of IL-22 levels, tissues were homogenized using a bead beater, and an ELISA for IL-22 (BioLegend) was performed according to the manufacturer’s instructions.

### Genomic comparisons of *Clostridium* species

Genomic DNA was isolated from overnight cultures of *C. immunis*, *C. symbiosum*, and *C. clostridioforme* using a MagAttract HMW DNA kit (Qiagen). A multiplexed sequencing library was generated using a SMRTbell prep kit (PacBio) and sequenced on a PacBio RS. Flye (release 2.8.3) was used to perform de novo assembly on quality-filtered reads^99^, which resulted in a complete and circularized genome for *C. immunis* (genome size of 5.34 Mbp), 108 contigs (genome size of 5.54 Mbp) for *C. symbiosum*, and 18 contigs (genome size of 5.84 Mbp) for *C. clostridioforme*. Gene annotation was performed with RAST^100^, and a genome-wide, sequence-based comparison of the three strains was performed using The SEED Viewer web server (version 2.0)^99^. Genes that were uniquely present in *C. immunis* were further manually curated to identify genes that are likely to generate extracellular products.

### EPS purification

The supernatant from overnight bacterial cultures was concentrated ∼50-fold using a 100 kDa molecular weight cutoff (MWCO) spin column (Amicon Ultra, Millipore). The retentate was incubated overnight with DNase I (50 μg/ml) and RNase A (50 μg/ml) at 37 °C followed by an overnight incubation with proteinase K (500 μg/ml) at 37 °C. The volume of the resultant mixture was increased to 15 ml with dH_2_O and concentrated back down to 1–2 ml using a 100 kDa MWCO column to remove proteinase K. The retentate was precipitated with ice-cold ethanol (80% final volume) at –20 °C overnight. Precipitates were air-dried and resuspended in distilled water.

### Visualization of EPS by gel electrophoresis

EPS (60 μg) was boiled for 5 mins in Laemmli buffer with β-mercaptoethanol and run on a 10% polyacrylamide gel (MiniPROTEAN TGX, BioRad) at 200 V for 1hr. Gels were visualized with a silver (Thermo Fisher Pierce), Sudan Black B (Thomas Scientific), or periodic acid–Schiff (PAS) stain. For PAS staining, the gel was washed in deionized water for 10 min, fixed in 12.5% trichloroacetic acid for 30 min, oxidized with 1% periodic acid for 1 h, washed in deionized water for 4–5 h, and stained with Schiff’s reagent for 1 h. After staining, the gel was reduced with three 10 min washes in 0.5% sodium metabisulfite. For the Sudan black B-stained gel, *E. coli* 055:B5 lipopolysaccharide (75 μg; Sigma) was used as a positive control. To assess the presence of nucleic acids, EPS (60ug) was run on a 1% agarose gel stained with a DNA Gel Stain (ApexBio) and imaged on an Odyssey XF system (LI-COR Biosciences).

### Glycosyl composition analysis

Glycosyl composition analysis was performed by combined gas chromatography-mass spectrometry (GC-MS) of the per-*O*-trimethylsilyl (TMS) derivatives of the monosaccharide methyl glycosides produced from the sample by acidic methanolysis as described previously^101^. Briefly, samples (150–300 μg) were heated with 1 M methanolic HCl in a sealed screw-top glass test tube for 18 h at 80 °C. After cooling and removal of the solvent under a stream of nitrogen, the samples were re-N-acetylated and dried again. The samples were then derivatized with Tri-Sil® (Pierce) at 80 °C for 30 min. GC-MS analysis of the TMS methyl glycosides was performed on an Agilent 7890A GC interfaced to a 5975C MSD, using a Supelco Equity-1 fused silica capillary column (30 m × 0.25 mm ID).

### Glycosyl linkage analysis

Glycosyl linkage analysis was performed as previously described^102^. Briefly, samples were dissolved in anhydrous DMSO overnight, and permethylation was achieved by two rounds of treatment with sodium hydroxide base (15 min) and iodomethane (25 min). The permethylated materials were hydrolyzed with 2 M TFA for 2 h at 121 °C and dried down with isopropanol under a stream of nitrogen. The samples were then reduced with 10 mg/mL NaBD_4_ in 100 mM NH_4_OH overnight, neutralized with glacial acetic acid, and dried with methanol. Finally, the samples were O-acetylated using 250 µL of acetic anhydride and 250 µL of concentrated trifluoroacetic acid (TFA) at 45 °C. The samples were dried under N_2_ stream, reconstituted in dichloromethane, and washed with nanopure water. The resulting partially methylated alditol acetates were analyzed on an Agilent 7890A GC interfaced to a 5975C MSD; separation was performed on a Supelco 2331 fused silica capillary column (30 m × 0.25 mm ID).

### 1H-NMR spectroscopy

Each lyophilized EPS sample (∼1–2 mg) was dissolved in 500 µl of D_2_O (99.9% D, Sigma) and placed in a 5-mm NMR tube. Sodium trimethylsilylpropanesulfonate (0.5 µl) was added as a reference. Liquid ^1^H-NMR data were obtained at 298 K on a Varian VNMRS spectrometer (1H, 599.66 MHz). ^1^H-NMR parameters: 2.0 s relaxation delay, 65536 Hz spectral width, 16384 data points and 64 transients with total recycle delay of 3.3 s between each transient. The experiment was performed with suppression of the HOD signal at 4.78 ppm by presaturation. Prior to the Fourier transformation, the data were apodized with an exponential decay function with line broadening of 0.5 Hz, 90⁰ sine square, and zero-filled to 64k points. The baselines were corrected automatically by subtracting a 3rd-order Bernstein polynomial fit. The spectra were processed and analyzed with MestreNova (version 14.2.1-27684).

### 2D NMR spectroscopy

EPS samples were dissolved in 1.0 ml of D_2_O (99.9 % D) and lyophilized. The samples were then dissolved in 0.5 ml of D_2_O with 2 µL of 50mM d6-DSS and transferred to 5-mm NMR tubes. All NMR data were acquired at 70 °C on a Bruker Avance III spectrometer (1H, 600.06 MHz) equipped with a cryoprobe using standard pulse sequences. A ^1^H spectrum was acquired using a standard pulse program with a spectral width of 9615.4 Hz, 32768 data points, and sixteen transients with a total recycle delay of 2 s. For the 2D HSQC experiments, 1H spectral widths were set to 72111.5 Hz and 13C spectral width to 10563.98 Hz. The 2D COSY spectrum was acquired with 8 transients per 400 increments. Chemical shifts were referenced to DSS (δH = 0.00 ppm and δC = 0.00 ppm). The spectra were processed and analyzed with MestreNova (version 14.0.1-23559).

### Slot immunoblot

EPS (5 μg) was added to a methanol-activated PVDF membrane housed in a slot blot manifold (Hoefer Inc.). The membrane was blocked in 3% bovine serum albumin in TBS containing 0.1% Tween 20 (TBS-T) for 1 h, followed by overnight incubation at 4 °C with an anti-phosphocholine antibody (1:500 dilution; clone BH8; Millipore). After washing with TBS-T, the membrane was incubated with an HRP-conjugated secondary antibody (Thermo Fisher Scientific) for 1 h. Chemiluminescence generated by treatment with Clarity ECL (Bio-Rad) was imaged on an Odyssey XF system (LI-COR Biosciences). EPS-treated membranes were also stained with PAS as detailed above to visualize the amount of carbohydrate loaded.

### Generation of *C. immunis*Δ*licABC*

We established that pMTL82151, one of the ClosTron plasmids^103^, fails to replicate in *C. immunis* and therefore could be used as a suicide vector. We used the Q5 High-Fidelity DNA polymerase (NEB) to PCR amplify the *ermB* gene that provides resistance to erythromycin (using pMTL83251 as the template) and two 800 bp regions (separated by 80 bp) that flanked the *licA* target site. PCR primers used are as follows: licA Left fwd: gctcggtacccggggatcctAATACCGTCAGCTACACTG; licA Left rev: cttcggccggTAGGCCGGACATATTCTATATC; licA Right fwd: gaatgtgtttTTTCTATATGGTGATCTTAGTAAAC; licA Right rev: agcttgcatgtctgcaggccTTTAATTGGACAAAGCGTC; ermB fwd: gtccggcctaCCGGCCGAAGCAAACTTAAG; ermB rev: catatagaaaAAACACATTCCCTTTAGTAACGTG. After digesting these three PCR products with XbaI and XhoI, they were cloned into an XbaI- and XhoI-digested pMTL82151 plasmid using HiFi assembly (NEB). Electrocompetent *E. coli* S17.Pir was transformed with the ligation mixture, which generated S17.Pir:pMTL82151ΔlicABC. This strain was used to transfer pMTL82151ΔlicABC into *C. immunis* using standard conjugation methods. Overnight cultures of *E. coli* donors and *C. immunis* recipients were mixed at a 10:1 donor-to-recipient ratio. Multiple aliquots (30 μl) of the mixture were spotted onto a BHI-S agar plate and incubated anaerobically at 37 °C. After 24–48 h, the total growth on each plate was collected, resuspended in fresh media, and grown on BHI-S plates containing erythromycin (500 μg/ml) and colistin (10 μg/ml). To confirm that the plasmid was not present in *C. immunis*, the culture was also grown on BHI-S plates containing colistin (10 μg/ml) and chloramphenicol (12.5 μg/ml), against which the plasmid backbone confers resistance. Erythromycin-resistant colonies of *C. immunis* were confirmed to harbor the insertion of the *ermB* gene and expected 80-bp deletion in *licA* by PCR and Sanger sequencing.

### Generation of *C. symbiosum:licABC*

We used the Q5 High-Fidelity DNA polymerase (NEB) to PCR amplify the *licABC* genes from *C. immunis*. PCR primers used are as follows: Ci licABC XbaI fwd: gatcctctagagtcgacAAACTCACTGGGGCGCTAGCAATG; Ci licABC XhoI rev: caggcctcgagatctccAATGCTAACAACCCTGACATCAGTG. After digesting this PCR product with XbaI and XhoI, it was cloned into an XbaI- and XhoI-digested pMTL83151 plasmid^103^. Electrocompetent *E. coli* S17.Pir was transformed with the ligation mixture, which generated S17.Pir::pMTL83151:licABC. This strain was used to transfer pMTL83151:licABC into *C. symbiosum* using standard conjugation methods detailed above.

### Cell culture

All tissues and cells were cultured in a humidified incubator with 5% CO_2_ at 37 °C. MNK-3 cells were obtained from Maria Ciofani (Duke University) and maintained as previously described^43^, adding recombinant mouse IL-7 (10 ng/ml; BioLegend) and IL-15 (10 ng/ml; BioLegend) every 3 days. After confirming no mycoplasma contamination, MNK-3 cells were authenticated by their ability to elicit the appropriate cytokine responses following stimulation as originally described^43^. Cells (5 x 10^5^) were plated in 24-well plates. The following day, cells were stimulated with IL-23 (10 ng/ml; BioLegend), IL-1β (10 ng/ml; BioLegend), IL-2 (10 ng/ml; BioLegend), phorbol myristate acetate (50 ng/ml; Sigma), ionomycin (500 ng/ml; Sigma), and with or without EPS (1.0 mg/ml) for 4 hours, after which they were lysed for RNA isolation.

### Flow cytometry

Single-cell suspensions of intestinal lamina propria cells were isolated and stained for flow cytometry as previously described^104^. Cells were stained with antibodies against CD3, CD19, CD11b, CD11c, Rorγt, and IL7Rα (all antibodies were from Biolegend). Data was acquired with a MacsQuant (Miltenyi), and analysis was performed with FlowJo software (BD).

### IL-22 depletion

Mice were injected intraperitoneally with either a neutralizing antibody against IL-22 (150 μg/dose; clone 8E11; Genentech) or an IgG1 isotype control (150 μg/dose) every other day. The antibody dosing regimen began 2 days prior to treatment with *C. immunis*.

### IL-22 administration

Mice were injected intraperitoneally with IL-22 (PeproTech) every other day. Animals received 50 ng/g body weight for the first dose and 25 ng/g body weight for all subsequent doses. For mice treated with *C. immunis* EPS, both treatments began the same day.

### FITC-dextran permeability assay

Mice were orally treated with or without *C. immunis*. One week later, mice were orally administered 0.8 g FITC-dextran (4 kDa; Sigma)/kg body weight. Four hours after FITC-dextran gavage, serum was collected. Fluorescence was measured in a plate reader (Biotek), and the concentration of FITC-dextran was determined based on a standard curve.

### Indirect calorimetry studies

We performed indirect calorimetry studies of *Ahr^fl/fl^* mice orally treated with or without *C. immunis* EPS as well as untreated ILC3-deficient littermate mice (*n*=8 male mice/group; ∼20 weeks of age). These studies were performed in Duke’s Mouse Behavioral and Neuroendocrine Analysis Core Facility. Mice were fed a high-fat diet (TestDiet Western Diet AIN-76A; 4.49 kcal/g; 40% of calories from fat) for 12 weeks, during which time they received weekly treatments with either sterile bacterial media (control) or *C. immunis* EPS. Mice were then placed into metabolic cages (Oxymax/CLAMS; Columbus Instruments), where they were monitored for 6 days. Data were analyzed in CalR^105^, and statistical analysis was performed using analysis of covariance (ANCOVA) testing and generalized linear models as implemented in CalR. Fecal samples were collected from the metabolic cages and examined by bomb calorimetry using a 6200 isoperibol bomb calorimeter (Pall) at the University of Michigan Animal Phenotyping Core.

### Respirometry

Respirometry experiments were performed in the Nutrition Obesity Research Center at the University of North Carolina, Chapel Hill using a XFe96 Analyzer (Seahorse Bioscience). Mice were treated with either sterile media (control) or 200 μg of *C. immunis* EPS. One week later, gonadal fat was isolated from the mice. Approximately 1.6 mg of minced gonadal fat was placed into 96-well spheroid microplates (Seahorse Bioscience), where the wells were coated with Cell-Tak (Corning) according to the manufacturer’s instructions. Each biological sample included 11–12 technical replicates. Tissue was incubated in pre-warmed XF Assay Media (modified DMEM with 7 mM glucose, 2 mM glutamine, 1 mM sodium pyruvate, pH 7.4; all reagents from Seahorse Bioscience) for 1 hr at 37°C in a CO_2_-free incubator. After a 10 min equilibration period, a XFe96 Analyzer (Seahorse Bioscience) was used to measure basal OCRs for 3 cycles, comprising a 3 min mix and a 2 min measure period. The OCRs were normalized to protein amounts present in the tissue lysate.

### Immunofluorescence and immunohistochemistry

Immunofluorescence and immunohistochemistry were performed as previously described^106^. Briefly, tissue slices from paraformaldehyde-fixed and paraffin-embedded tissue blocks were dewaxed and dehydrated using xylene and a graded ethanol series. Antigen retrieval was performed by boiling the slides in 10 mM sodium citrate buffer (pH 6.0) for 30 min. Slides were treated with hydrogen peroxide for 10 min to block endogenous peroxidases, after which the slides were incubated with a blocking serum for 10 min at room temperature. For immunofluorescence detection of NFIL3, slides were incubated with a polyclonal rabbit anti-NFIL3 antibody (1:400; Proteintech) overnight at 4 °C, followed by an AlexaFluor 555-conjugated goat anti-rabbit IgG secondary antibody (1:200; ThermoFisher) for 2 h at room temperature. Slides were counterstained with DAPI (Sigma) for nuclear visualization, and images were acquired using a LSM 880 Airyscan fast inverted confocal microscope (Zeiss). NFIL3 fluorescence was quantified using ImageJ software. For immunohistochemistry detection of UCP1, slides were incubated with a polyclonal rabbit anti-UCP1 antibody (1:1000, Abcam) overnight at 4 °C. The following day, antibody reactivity was detected using a rabbit specific HRP/DBA detection IHC kit (Abcam), following the manufacturer’s instructions. Between each incubation step, slides were washed with PBS four times for 3 min each. Slides were counterstained with hematoxylin, and the slides were imaged using an Axio Imager Z2 Widefield Fluorescence Microscope (Zeiss)

### Cultures of stromal vascular fractions (SVF) from adipose tissue

Isolation of SVF and induction of beige cells were performed as previously described^53^. Briefly, iWAT and gWAT from C57BL/6 mice was digested with collagenase D (1 mg/ml; Sigma) at 37°C for 40 min with shaking. The cell suspension was filtered through a 100 μm cell strainer and centrifuged at 500 x *g* for 5 min at 4 °C to separate floating adipocytes from the pelleted SVF, which was resuspended in DMEM/F12 containing 10% FBS. This SVF fraction was then treated with RBC lysis buffer (Chemcruz) for 5 min at room temperature, passed through a 100 μm filter, and plated in either 6 cm dishes or 24-well plates. Cells were grown to ∼95% confluence and then differentiated into beige adipocytes as previously described^107^. Briefly, beige cell induction was conducted by culturing cells for 2 days in media containing 10% FBS, 0.5 mM isobutylmethylxanthine, 125 nM indomethacin, 5 μM dexamethasone, 850 nM insulin, 1 nM T_3_, and 1 μM rosiglitazone, after which media was changed to contain 10% FBS, 850 nM insulin, 1 nM T_3_, and 1 μM rosiglitazone for ongoing maintenance. IL-22 (100 ng/ml; PeproTech) was added to the induction and maintenance media for some samples. Six days later, cells were either lysed for RNA isolation or processed for immunofluorescence. For the latter, wells were incubated with a polyclonal rabbit anti-UCP1 antibody (1:1000, Abcam) followed by an AlexaFluor 555-conjugated goat anti-rabbit IgG secondary antibody (1:200; ThermoFisher). Cells were counterstained with DAPI (Sigma) for nuclear visualization, and images were acquired using a LSM 880 Airyscan fast inverted confocal microscope (Zeiss). ImageJ software was used to quantify UCP1 fluorescence intensity and total cell number. All chemicals were obtained from Sigma unless indicated otherwise.

### Phylogeny of *C. immunis licABC* homologs

Homologs of the *C. immunis licABC* locus were identified using BLASTn (2.13.0). The gene sequences for these *licABC* loci as well as the sequences from select organisms primarily found in the respiratory tract were aligned using ClustalW. A maximum likelihood phylogenetic tree was generated using RAxML as implemented with the default settings in MegAlign Pro (DNA Star).

### Meta-analysis of *licC* abundances in human studies

We obtained publicly available human-derived shotgun metagenomic data from the curatedMetagenomicData R package (version 3.6.2)^108^. We included all studies available as of July 2023 that had stool samples from westernized countries for which there were corresponding BMI and/or serum triglyceride data. We required samples to be labeled as “control” for “study condition” and “healthy” for “disease.” To ensure independence in the dataset (i.e., having only one sample per subject), we included only the baseline sample (or the first entry when no timepoint information was provided) for studies with multiple samples per subject. Additionally, for individuals included in multiple studies, we only included their data from one study. For the BMI analysis, we defined groups as heathy (BMI < 25 kg/m^2^), overweight (25 kg/m^2^ ≤ BMI < 30 kg/m^2^), or obese (BMI ≥ 30 kg/m^2^), and we required studies to have ≥10 obese or non-obese subjects, which resulted in a total of 11 studies (total of 4,145 individuals) being combined for this meta-analysis^109–119^. For the serum triglyceride analysis, we defined groups as normal (triglyceride < 150 mg/dL), borderline (150 mg/dL ≤ triglyceride < 200 mg/dL), or high (triglyceride ≥ 200 mg/dL), and we required studies to have ≥10 subjects, which resulted in a total of 4 studies (total of 247 individuals) being combined for this meta-analysis^113,120–122^.

We downloaded HUMAnN3-processed UniRef90 gene family abundance tables for each of these studies and scaled them to copies per million. To identify homologs of the *licC* gene, we used the *C. immunis licC* gene sequence to search for UniRef90 entries in the UniProt Knowledgebase^123^, setting a similarity threshold of ≥40%. We used the grep UNIX command to extract the abundance of these *licC* homologs from the study-associated data, and we summed their abundances.

### Statistics

Sample-size estimates for each experiment were based on previous laboratory experience. The investigators were not blinded to allocation during experiments and outcome assessment. Prism 10 (GraphPad Software) was used for all statistical analyses unless otherwise specified, and the specific statistical tests used are included in the figure legends.

### Data Availability

The genome sequences for *C. immunis*, *C. symbiosum*, and *C. clostridioforme* have been deposited at GenBank under accession number xxxxx. The RNAseq data have been deposited to NCBI under BioProject accession number xxxxx.

