## Supplementary figures and images for "A commensal-derived sugar protects against obesity by regulating immunometabolism"

### Supplementary Figures

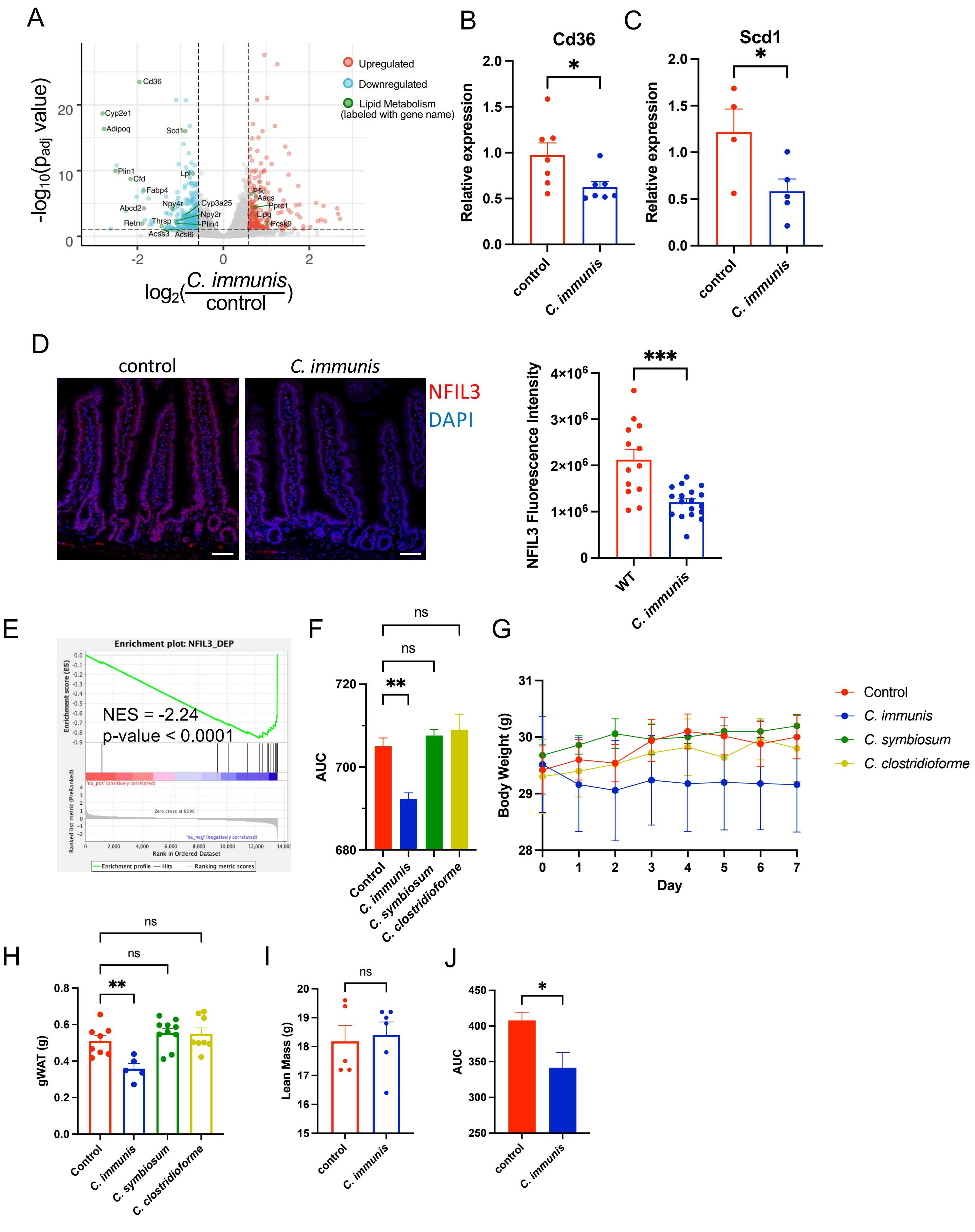

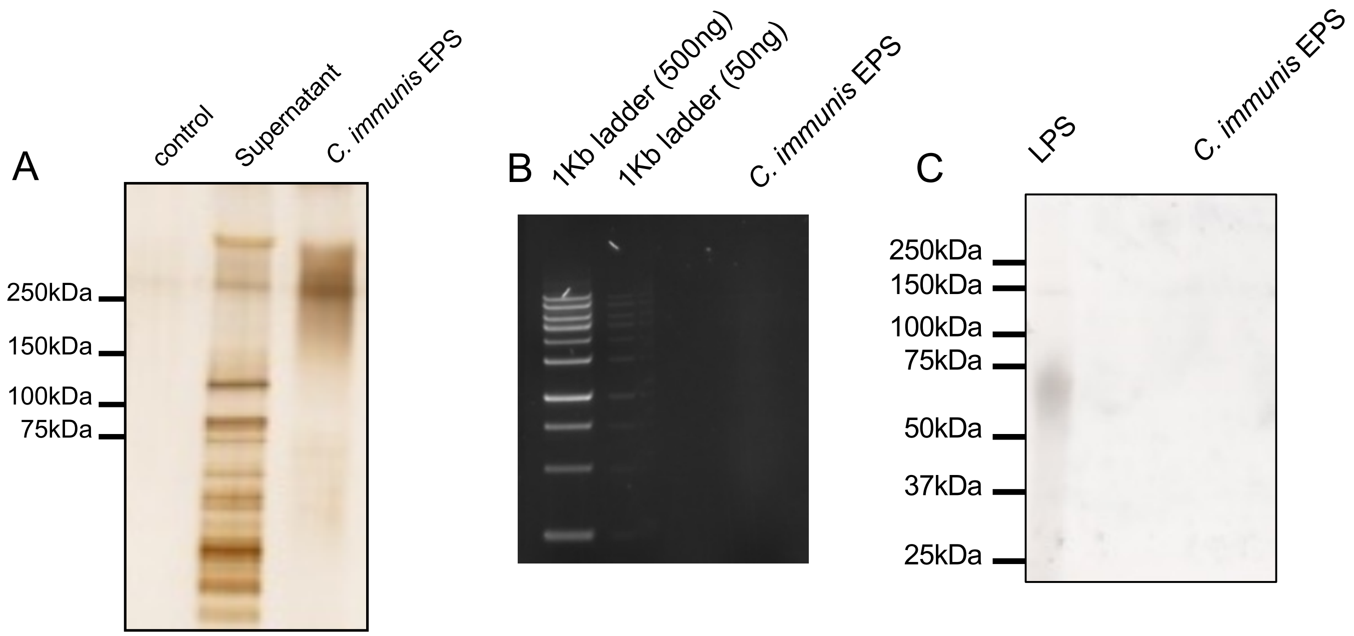

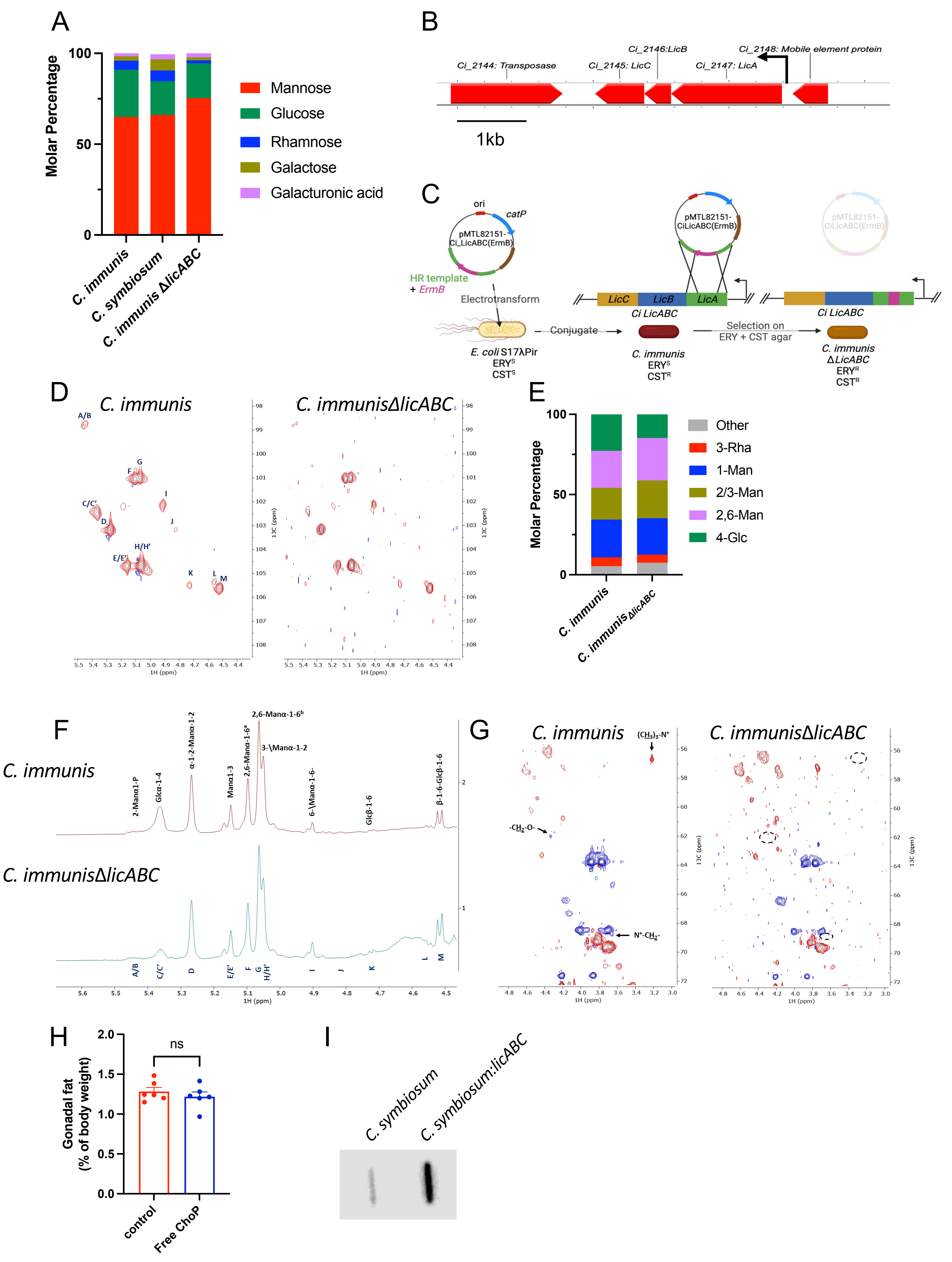

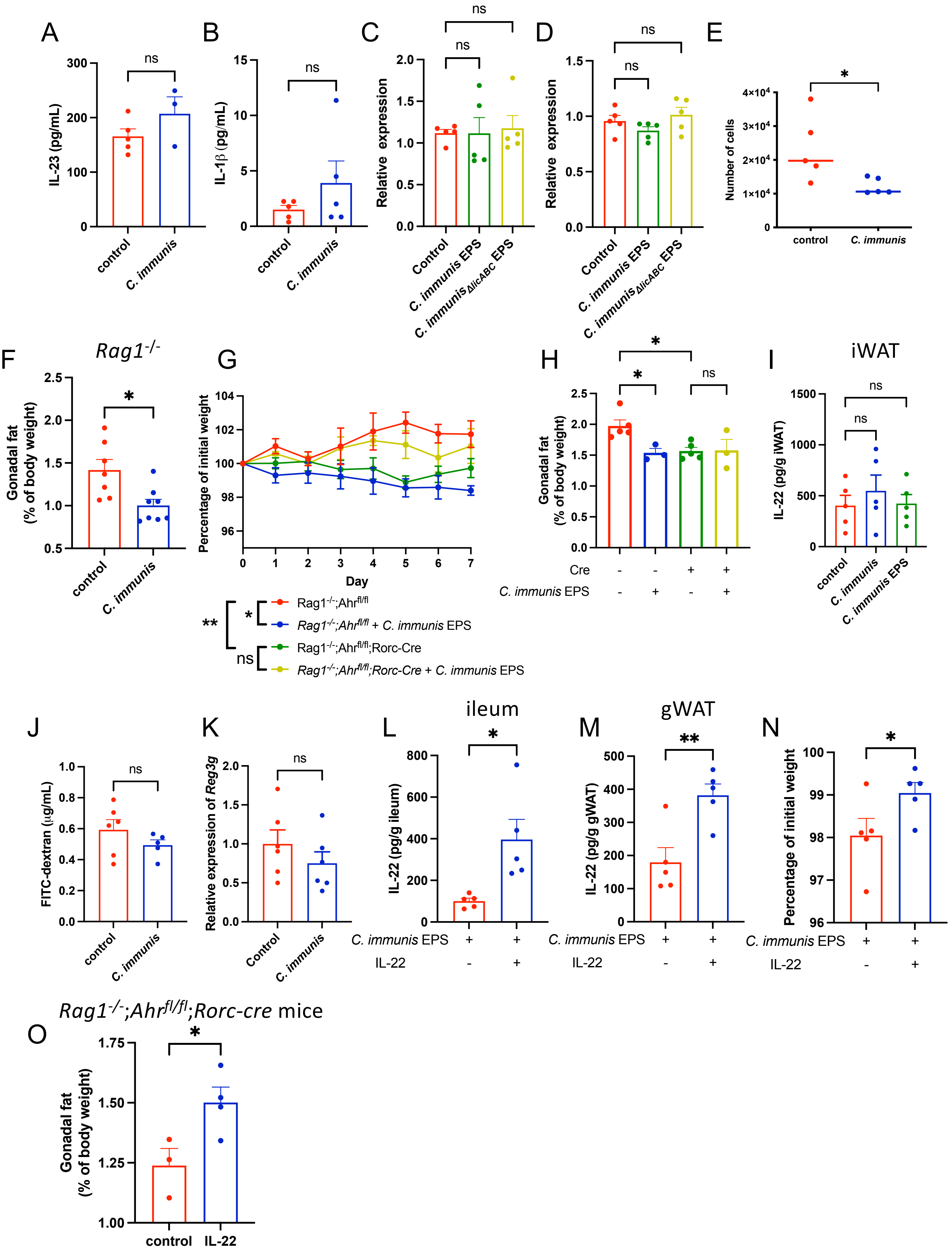

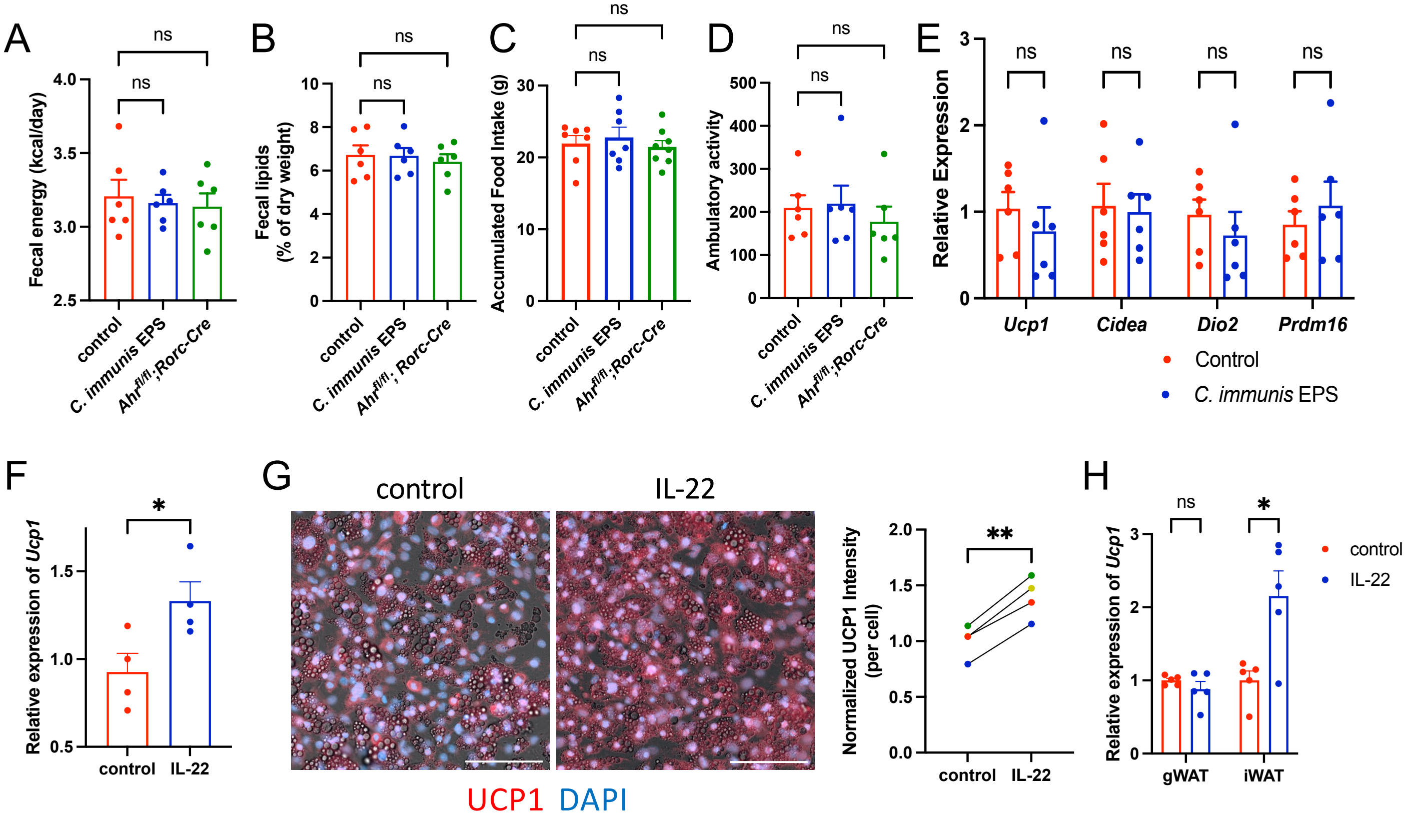

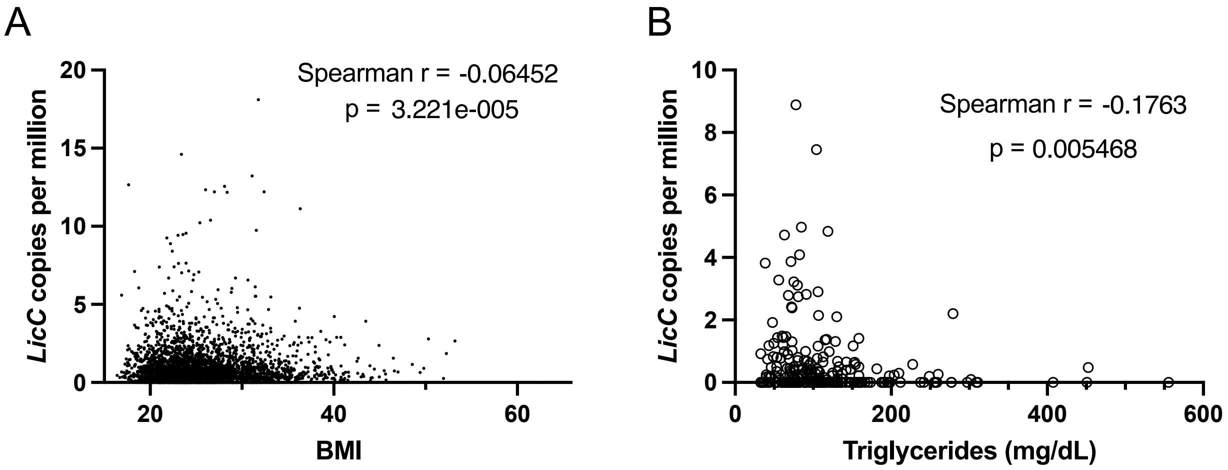
