## Supplementary Table 1 for "A commensal-derived sugar protects against obesity by regulating immunometabolism"

**Table S1. *C. immunis* genes identified by comparative genomics and predicted to impact extracellular products**. Bolded genes are related to polysaccharide biosynthesis.

| Gene ID | Function |
|---|---|
| Ci_66 | Tripartite tricarboxylate transporter TctA family |
| Ci_131 | NAD-dependent oxidoreductase |
| Ci_271 | Protein-export membrane protein SecD/SecF |
| Ci_554 | Hydrolase of alpha/beta superfamily |
| **Ci_818** | **capsular polysaccharide biosynthesis protein Cps4F** |
| **Ci_820** | **Capsular polysaccharide synthesis enzyme Cap5F** |
| Ci_821 | UDP-N-acetyl-L-fucosamine synthase |
| Ci_891 | Pantothenate:Na+ symporter |
| Ci_1692 | Branched-chain amino acid transport system permease protein LivM |
| **Ci_2065** | **capsular polysaccharide biosynthesis protein** |
| **Ci_2145** | **Pyrophosphorylase involved in lipopolysaccharide biosynthesis (LicC)** |
| Ci_**2146** | **Capsular polysaccharide biosynthesis protein (LicB)** |
| Ci_2162 | Acyltransferase 3 |
| Ci_2655 | Oligopeptide ABC transporter, periplasmic oligopeptide-binding protein OppA |
| Ci_2783 | Multi antimicrobial extrusion protein (Na(+)/drug antiporter) |
| Ci_2978 | PTS system, mannitol-specific component |
| Ci_3330 | Glycosyltransferase, group 2 family |
| Ci_3460 | Cell surface protein |
| Ci_3691 | Glycoside-Pentoside-Hexuronide transporter |
| Ci_3700 | UDP-N-acetyl-L-fucosamine synthase |
| **Ci_3701** | **Capsular polysaccharide synthesis enzyme Cap5F** |
| Ci_3710 | Galacturonosyl transferase |
| Ci_3711 | Alpha-1,3-N-acetylgalactosamine transferase PglA |
| Ci_3992 | Type IV fimbrial assembly protein PilC |
| Ci_4441 | Sodium/glutamate symporter |
| Ci_4572 | Bacteriocin-like protein |
| Ci_5479 | Polygalacturonase |
| **Ci_5817** | **Putative enzyme of poly-gamma-glutamate biosynthesis (capsule formation)** |
| Ci_5925 | Choline binding protein A |
| Ci_5927 | Choline binding protein A |
